# A Randomized, Controlled Clinical Trial Demonstrates Improved Cognitive Function in Senior Dogs Supplemented with a Senolytic and NAD+ Precursor Combination

**DOI:** 10.1101/2024.02.26.581616

**Authors:** Katherine E. Simon, Katharine Russell, Alejandra Mondino, Chin-Chieh Yang, Beth C Case, Zachary Anderson, Christine Whitley, Emily Griffith, Margaret E. Gruen, Natasha J. Olby

## Abstract

Age-related decline in mobility and cognition are associated with cellular senescence and NAD+ depletion in dogs and people. A combination of a novel NAD+ precursor and senolytic, LY-D6/2 was examined in this randomized controlled trial. Seventy dogs were enrolled and allocated into placebo, low or full dose groups. Primary outcomes were change in cognitive impairment measured with the owner-reported Canine Cognitive Dysfunction Rating (CCDR) scale and change in activity measured with physical activity monitors. Fifty-nine dogs completed evaluations at the three-month primary endpoint, and 51 reached the six-month secondary endpoint. There was a significant difference in CCDR score across treatment groups from baseline to the primary endpoint (p=0.02) with the largest decrease in the full dose group. There were no significant differences between groups in changes in measured activity. However, the proportion of dogs that improved in frailty and owner-reported activity levels and happiness was higher in the full dose group than other groups. Adverse events occurred equally across groups. All groups showed improvement in cognition, frailty, and activity suggesting placebo effect and benefits of trial participation. We conclude that LY-D6/2 significantly improves owner-assessed cognitive function and may have broader effects on frailty, activity and happiness as reported by owners.

## Introduction

Improved veterinary medical care and migration of dogs from a working role to family member have resulted in extension of the canine lifespan [1,2,3]. As a result, dogs, like humans, experience a wide range of age-related morbidities [4,5,6,7,8,9]. Indeed, inclusion of dogs in the family household with exposure to the same environmental contaminants and stressors, and adoption of the activity patterns and sometimes nutrition of their owners has resulted in a very similar array of these conditions occurring in both species [10,11].

Cognitive function and mobility have been proposed as key hallmarks of functional aging, and their age-associated attrition appears to be linked in both humans [12,13,14] and dogs [15,16]. Normal age-related cognitive changes include alterations in sleep [17], memory [18,19], attention [20,21] and social interactions [22,23]. In a high proportion of people and dogs, these develop into Alzheimer’s Disease [24] or Canine Cognitive Dysfunction Syndrome (CCDS) [25,26,27,28] respectively. Similarly, activity and mobility decrease with age in humans [29] and dogs [30] due to sarcopenia [31], decreased motivation [32] and common pathologies such as osteoarthritis [33]. These changes affect the patient’s quality of life, and take a significant toll on the caregiver [34,35,36]. Supplements that can reduce cognitive decline and maintain mobility in aging dogs would have enormous impact on dogs and their caregivers, and translational relevance for aging people.

Clinical manifestations of aging have their origin at a molecular level with 12 molecular hallmarks of aging now recognized [37]. The burgeoning field of geroscience has identified therapeutic targets to mitigate the aging process within these hallmarks and numerous clinical trials are now underway in people [38]. While the human anti-aging field is flourishing, far fewer studies target age-related decline in dogs. In this randomized, controlled, double-blinded clinical trial we evaluated a combination of supplements that target two important hallmarks of aging, cellular senescence and depletion of cellular nicotinamide adenine dinucleotide (NAD+) concentrations.

Senolytics have emerged as a promising anti-aging therapeutic strategy [39,40]. Cellular senescence increases with age and while it can be a protective mechanism, it can also cause an unwanted inflammatory response, the Senescence-Associated Secretory Phenotype (SASP) [37,41]. Senolytics include plant derived flavonoids such as quercetin and fisetin that act through inhibition of anti-apoptotic proteins (such as BCL-2 family proteins) [42]. Reducing senescence reduces age-related pathology *in vitro* and extends lifespan and a variety of different functional outcomes *in vivo* [43,44]. Senolytics are marketed widely as anti-aging supplements and human clinical trials evaluating senolytics in age-related diseases such as osteoarthritis, heart disease and Alzheimer’s disease are underway [45], but clinical trials in dogs are lacking.

Supplementation with an NAD+ precursor takes a more broadly targeted approach to aging. NAD+ is involved in regulation of key biological processes, cellular signaling and electron transfer, but it declines with age [41,46,47], exacerbating age-related diseases [48,49]. Precursors such as nicotinamide mononucleotide have been shown to mitigate the effects of aging and age-related pathologies [50] and restoring levels of NAD+ improves cellular energy and metabolism, leading to improvements in health and lifespan [51,52]. Human clinical trials are ongoing [53], however there is only one canine study published to date, where muscle function improved following treatment with nicotinamide in dogs with Duchenne’s Muscular Dystrophy [54]. Senescent cells secrete an NADase (CD38), causing depletion of NAD+. As a result, it has been suggested that combining NAD+ precursors with a senolytic amplifies NAD+ anti-aging properties [55].

This study evaluated the effects of a repeated monthly regimen of two consecutive days of the senolytic (LY-D6^TM^) and NAD precursor followed by daily NAD precursor (LY-D2^TM^) on cognition and activity in companion dogs. We hypothesized that supplementation with LY-D6/2 combination would reduce cognitive and activity level decline in aging dogs. Companion dogs aged 10 years or older were randomized to receive either placebo or LY-D6/2 combination at two different doses (low and full) with primary outcomes of change in owner assessed cognition and collar mounted physical activity monitor (PAM) assessed activity levels after 3 months.

## Results

Two hundred and forty-three surveys were completed, 91 dogs were brought in for screening appointments from which 70 dogs were enrolled and 67 completed baseline assessments (Figure 1). One dog was withdrawn due to pre-existing disease and 66 completed the one-month assessments. Two dogs died, one withdrew and one was excluded due to noncompliance, leaving 62 dogs to complete the three-month (primary endpoint) assessments. Four dogs were euthanized between month three and six and two withdrew due to mobility decline and neck pain. Fifty-six dogs completed all visits. Two dogs had active urinary tract infections (UTIs) at their six-month visit so their data were excluded from the six-month analyses. Data from three dogs were excluded from three- and six-month analyses due to treatment noncompliance (>20% missed doses).

**Figure 1:**
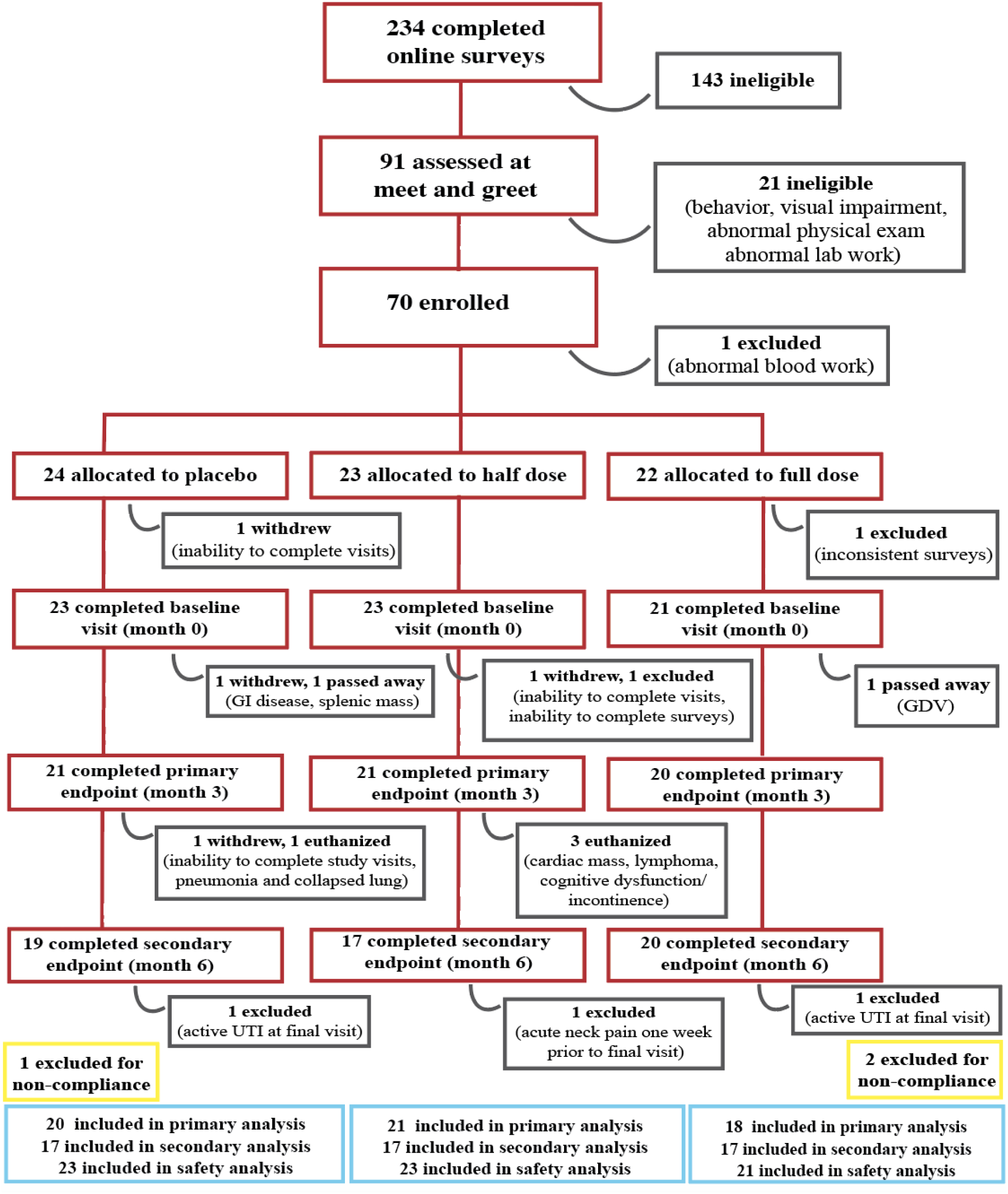
Participant Flowchart. Diagram featuring patient screening, treatment allocation, attrition, and exclusion throughout the trial. GDV-Gastric Dilation-Volvulus. UTI-Urinary Tract Infection.

### Study Population

Demographic details and baseline outcome measures of dogs at time of enrollment are provided in Table 1. The only significant difference between groups in baseline characteristics was age (p=0.02) with the full dose group being significantly older than the placebo or low dose groups. Owners were asked to maintain consistency in their pet’s routine over the course of the trial, but the number of pets changed in 14 households (addition or euthanasia of a pet).

**Table 1:** Population demographics and outcome measures of study participants at enrollment by group.

|  | Placebo (n=23) | Low Dose (n=23) | Full Dose (n=21) | P value | Fisher's Exact Test |
| --- | --- | --- | --- | --- | --- |
| Age (years) | Mean: 12.94<br>St Dev: 1.45 | Mean: 12.25<br>St Dev: 1.42 | Mean: 13.63<br>St Dev: 1.29 | 0.01 |  |
| Fractional Lifespan | Median: 1.10<br>Range: 0.76-1.24 | Median: 1.03<br>Range: 0.72-1.22 | Median: 1.12<br>Range: 0.87-1.23 | 0.19 |  |
| Sex | Female: 10 (43.5%)<br>Male: 13 (56.5%) | Female: 9 (39.1%)<br>Male: 14 (60.9%) | Female: 12 (57.1%)<br>Male: 9 (42.9%) | 0.393 |  |
| Weight (kg) | Median: 24.5<br>Range: 7.9-39.5 | Median: 27.7<br>Range: 6.5-45 | Median: 22.9<br>Range: 8.3-33.2 | 0.25 |  |
| Body Condition Score | 3/9: 1 (4.3%)<br>4/9: 3 (13.0%)<br>5/9: 13 (56.5%)<br>6/9: 3 (13.0%)<br>7/9: 3 (13.0%) | 4/9: 2 (8.7%)<br>5/9: 13 (56.5%)<br>6/9: 8 (34.8%) | 4/9: 5 (23.8%)<br>5/9: 12 (57.1%)<br>6/9: 2 (9.5%)<br>7/9: 1 (4.8%)<br>8/9: 1 (4.8%) | 0.15 | 0.19 |
| Frailty Status | Non-frail: 15 (65.2%)<br>Frail: 8 (34.8%) | Non-frail: 12 (52.2%)<br>Frail: 11 (47.8%) | Non-frail: 11 (52.4%)<br>Frail: 10 (47.6%) | 0.59 |  |
| Frailty Score | 1/5: 6 (26.1%)<br>2/5: 9 (39.1%)<br>3/5: 4 (17.4%)<br>4/5: 3 (13.0%)<br>5/5: 1 (4.3%) | 0/5: 1 (4.3%)<br>1/5: 5 (21.7%)<br>2/5: 6 (26.1%)<br>3/5: 8 (34.8%)<br>4/5: 1 (4.3%)<br>5/5: 2 (8.7%) | 1/5: 3 (14.3%)<br>2/5: 8 (38.1%)<br>3/5: 9 (42.9%)<br>4/5: 1 (4.8%) | 0.42 | 0.56 |
| Pain Score | Median: 7<br>Range: 0-20 | Median: 5<br>Range: 1-15 | Median: 6<br>Range: 2-21 | 0.61 |  |
| CCDR Score | Median: 38<br>Range: 34-46 | Median: 40<br>Range: 34-57 | Median: 39<br>Range: 34-58 | 0.28 |  |
| Inhibitory Control (Cylinder Task) | Median: 87.5<br>Range: 0-100 | Median: 87.5<br>Range: 0-100 | Median: 75<br>Range: 0-100 | 0.57 |  |
| Detour | Median: 37.5<br>Range: 0-87.5 | Median: 25<br>Range: 0-100 | Median: 12.5<br>Range: 0-100 | 0.52 |  |
| Sustained Gaze (sec) | Median: 22.78<br>Range: 5.92-60 | Median: 17.74<br>Range: 4.28-55.56 | Median: 15.09<br>Range: 2.12-60 | 0.22 |  |
| Off Leash Gait Speed (m/s) | Median: 1.51<br>Range: 0.54-3.56 | Median: 1.39<br>Range: 0.59-3.03 | Median: 1.44<br>Range: 0.66-3.81 | 0.55 |  |

Twenty-eight dogs had a change in medication; we compared the number of household and medication changes from baseline to primary endpoint (month three) and from primary endpoint to secondary endpoint (month six) across groups and found no significant differences between groups (Supplementary Table S1).

## Primary Outcomes

### CCDR

#### Primary Endpoint (3 months)

All dogs entered the trial with mild to moderate cognitive impairment. The CCDR scores did not differ significantly between groups (p=0.28) (Table 1). Over the first three months of the trial, CCDR score decreased (improved) in each treatment group. Individual frailty status at enrollment was significantly associated with change in CCDR and so was incorporated into a repeated measures model comparing CCDR score between groups at baseline, one and three months. There was a significant difference between treatment groups over the three-month period (p=0.02), with the full dose group showing the largest decrease (improvement) in CCDR score (Figure 2).

**Figure 2:**
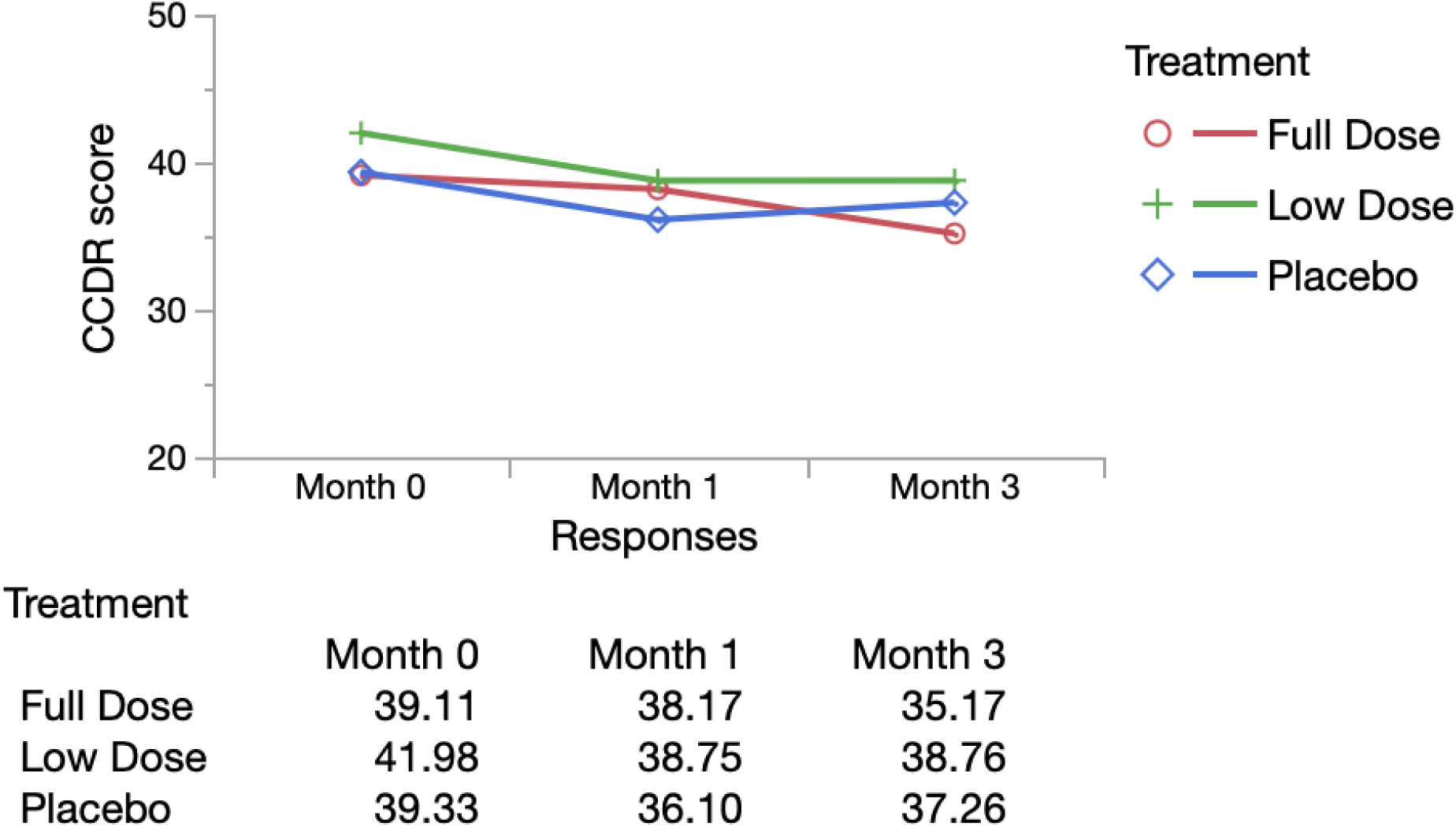
Repeated Measures Analysis of CCDR Score (Adjusted for Baseline Frailty Status) Mean CCDR scores by group adjusted for individual baseline frailty status over time from repeated measures model (MANOVA analysis in JMP). Wilks’ lamda value was evaluated, with p<0.05 indicating a significant difference between groups. All original outcome measure data is provided in Supplementary Table S5.

Change in CCDR score was also categorized as failure (increase in score representing worsening cognition) or success (static or decreased score) and groups were compared using a chi-square analysis. In the full dose group 16/18 (88.9%) dogs were successes, compared with 15/21 (71.34%) of low dose dogs and 12/20 (60%) of placebo dogs. However, these differences were not statistically significantly (p=0.11).

#### Secondary Endpoint (6 months)

We next assessed whether individuals maintained their CCDR scores to the final endpoint. Score differences from month three to month six were calculated and compared across groups. Median change in CCDR was static in all groups (Supplementary Table S2), with no significant difference across groups (p=0.44).

### Activity Monitor

Activity was initially analyzed using functional linear modeling (FLM) to allow patterns of change over the entire 24-hour period to be captured. At baseline, activity levels peaked around 6-10am and 4-9pm, reflecting times when owners interact with their dog. There was no significant difference between groups (Figure 3a,b). Weekend activity did differ between groups around 12pm, reaching the global threshold for significance, with the full dose group showing higher activity (Figure 3c,d).

**Figure 3:**
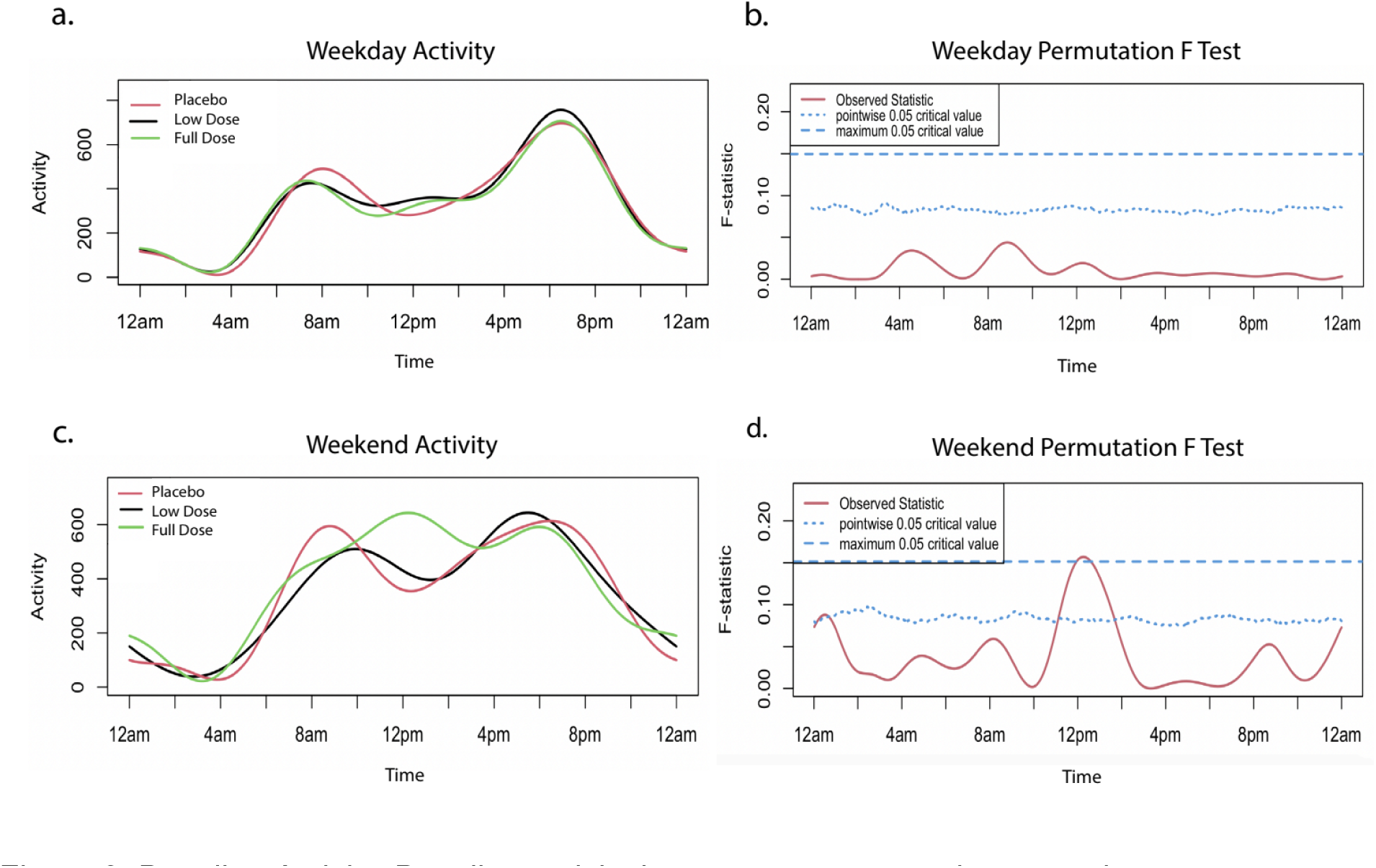
Baseline Activity. Baseline activity by treatment group prior to starting treatment on weekdays and weekends. The graphs (a,c) on the left illustrate average weekday/weekend activity levels over a 24-hour period. The graphs on the right (b,d) indicate the level of significance between groups. When the observed statistic (red line) is above the dashed and dotted lines (blue line), it indicates a global or pointwise significant difference between groups respectively. No significant differences were observed during weekdays. A global threshold for significance is reached around 12pm on weekends. A pointwise significant difference is observed around 12am on weekends. All original activity data is provided in Supplementary Table S10.

#### Primary Endpoint

When change in activity level from baseline to month three was examined on weekdays, groups maintained similar levels of activity compared to baseline. Notably, all groups showed a small increase in morning and evening activity and decrease in afternoon activity with no significant difference between them (Figure 4a,b). Changes in weekend activity were more variable, reflecting the owners’ less defined weekend schedules (Figure 4c). The placebo and full dose groups showed larger fluctuations than the low dose group, but the differences between groups did not reach global significance (Figure 4c,d).

**Figure 4:**
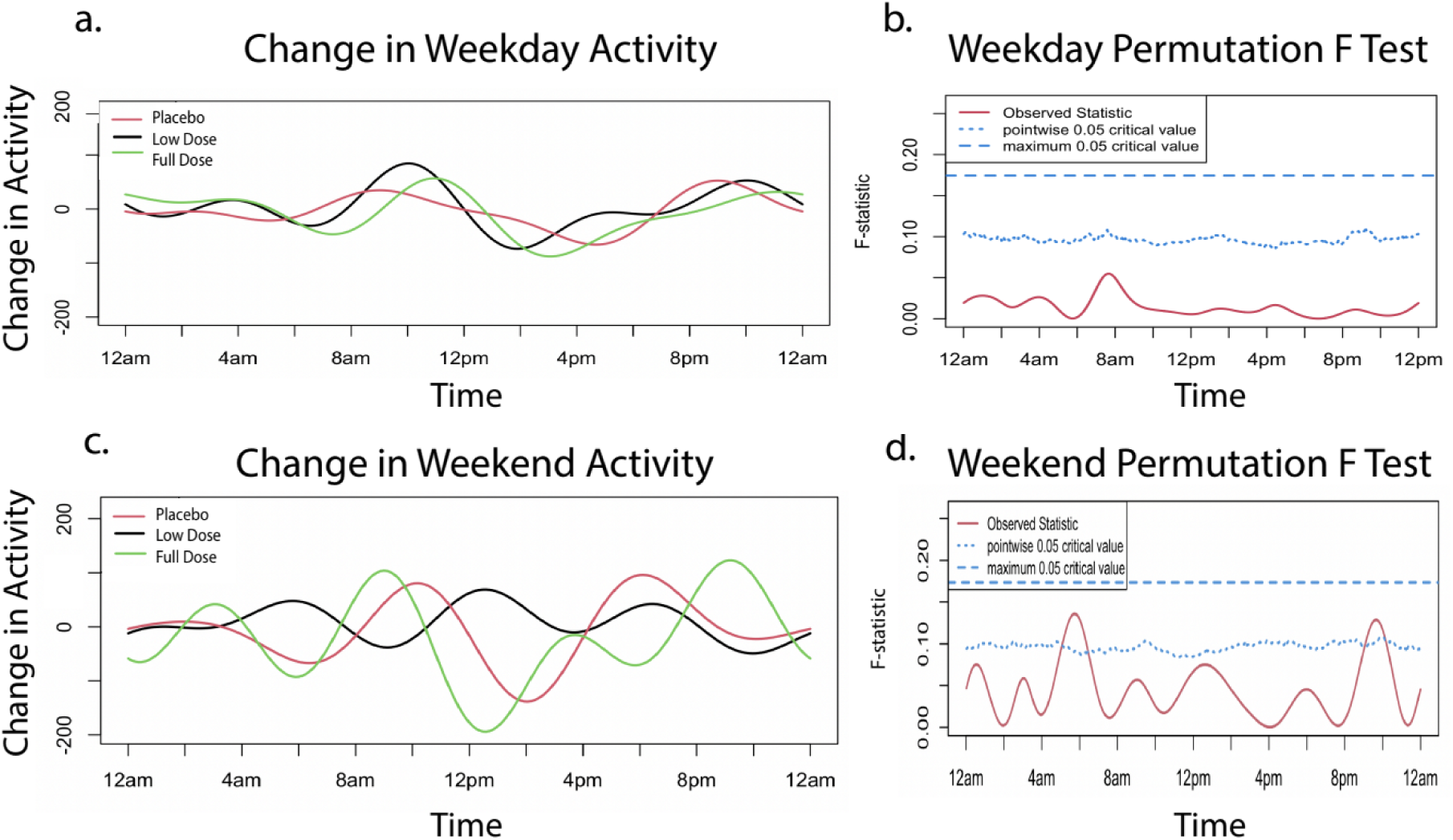
Change in activity from month 0 to month 3. Change in activity from baseline to primary endpoint on weekdays and weekends by treatment group. Activity at baseline was subtracted from activity at month three to obtain the change in activity utilized in the FLM analysis. The graphs on the left (a,c) illustrate change in weekday/weekend activity levels over a 24-hour period. The graphs on the right (b,d) indicate the level of significance between groups. When the observed statistic (red line) is above the dashed and dotted blue lines, it indicates a global or pointwise significant difference between groups respectively. There was no significant difference between group activity over the weekdays. On the weekends, pointwise significance was reached at approximately 6am and 10pm. All original activity data is provided in Supplementary Table S10.

#### Secondary endpoint

Similarly, when evaluating the change in activity from months three to six, activity remained relatively unchanged on weekdays, apart from an increase in all groups in the afternoon to evening, while the weekend activity fluctuated more widely (Figure 5a,c). There was no significant difference between groups on weekdays but at the weekend, groups differed significantly around 6am with the low dose groups showing a decrease in activity (Figure 5d).

**Figure 5:**
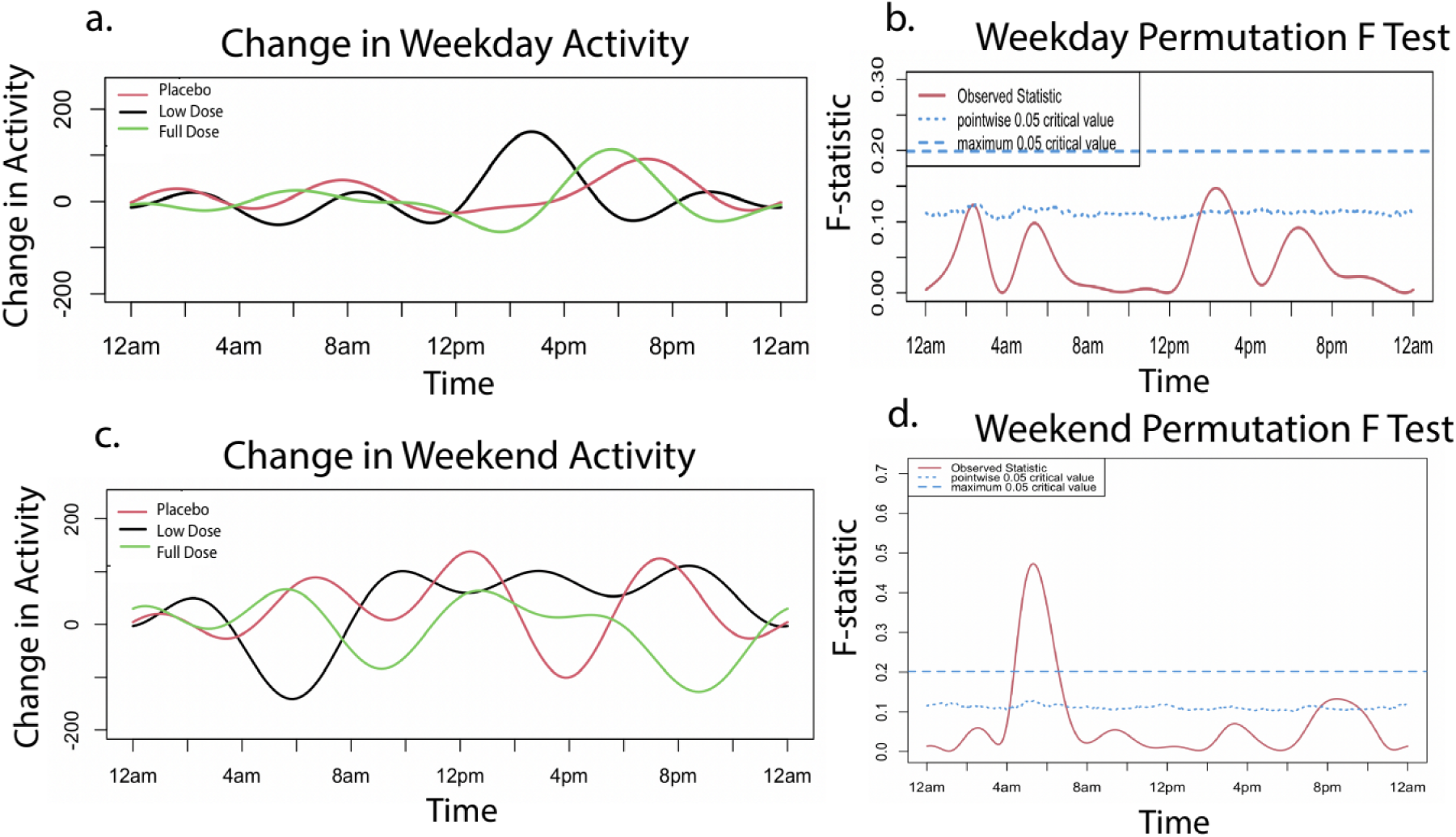
Change in activity from month 3 to month 6. Change activity from primary endpoint (month three) to secondary endpoint (month six) on weekdays and weekends by treatment group. Activity at month three was subtracted from activity at month six to obtain the change in activity utilized in the FLM analysis. The graphs on the left illustrate average change in weekday/weekend activity levels over a 24-hour period. The graphs on the right indicate the level of significance between groups. When the observed statistic (red line) is above the dashed and dotted lines (blue line), it indicates a global or pointwise significant difference between groups respectively. Pointwise significance is observed to reach threshold level on weekdays around 2am and 2pm. Weekend activity reaches the global significance level at 6am as well as a pointwise significance level around 9pm. All original activity data is provided in Supplementary Table S10.

#### Cumulative Activity

To incorporate covariates into the analysis, total cumulative activity for each dog was examined using repeated measures models. Weekday daytime activity was affected by age, and weekday and weekend daytime activity were affected by sex. Neither weekday nor weekend nights required correction for covariates. The repeated measures models for summated activity showed no significant difference over time (from baseline to primary endpoint) between groups during the day (weekday: p=0.95, weekend: p=0.71) or during the night (weekday: p=0.31, weekend: p=0.65) (Supplementary Figure S3). When assessing the difference in cumulative activity between three and six months, we found no significant difference across either time period on weekdays (day: p=0.89, night: p=0.10) or weekends (day: p=0.60, night: p=0.83)(Supplementary Table S4).

#### Owner assessed Activity

By contrast, when owners were asked to classify activity as static, reduced or increased at the primary endpoint, 8/18 (44%) owners in the full dose group reported static and 7/18 (39%) reported increased activity levels, compared with 13/21 (62%) static and 2/21 (10%) increased in the low dose group and 11/20 (55%) static and 4/20 (20%) increased in the placebo group (Supplementary Table S5). This difference between the groups was not significant (p=0.29). A majority of dogs across all groups remained static at the final endpoint: 10/17 (59%) of the full dose group, 12/17 (71%) of the low dose group and 11/17 (65%) of the placebo group. The differences were not significant (p=0.64).

## Secondary Outcomes

### Primary Endpoint

#### Frailty Score

The proportion of dogs that were frail (3 or more of 5 domains classed as impaired) and the number of impaired domains at study outset for each treatment group are provided in Table 1; groups were not significantly different. As expected with a senior pet population, there were changes in frailty over the course of the study which were categorized as success (static or improved) or failure (deteriorated) (Supplementary Table S5). The majority of dogs in full dose (13/18, 72.2%) and low dose (16/21, 76.2%) groups were classified as success after 3 months, as compared with 11/20 dogs (55%) in the placebo group; however groups did not differ significantly (p=0.32).

#### Cylinder Task (Inhibitory control)

Inhibitory control is a test of executive function, similar to impulse control, which decreases with age in both humans and dogs [56,57]. The full dose group started with a lower mean baseline score (66.1%) than the low dose (78.79%) and placebo group (76.6%); this difference was not significant (p=0.57) (Table 1). When assessing change from baseline to the primary endpoint, inhibitory control required correction for changes to household and changes to medication. These were therefore incorporated into the repeated measures model, which showed all groups increasing in score (improving their performance in the task) (Figure 6a). However, all groups showed a similar trajectory of improvement and there was no significant difference over time across groups (p =0.95).

**Figure 6:**
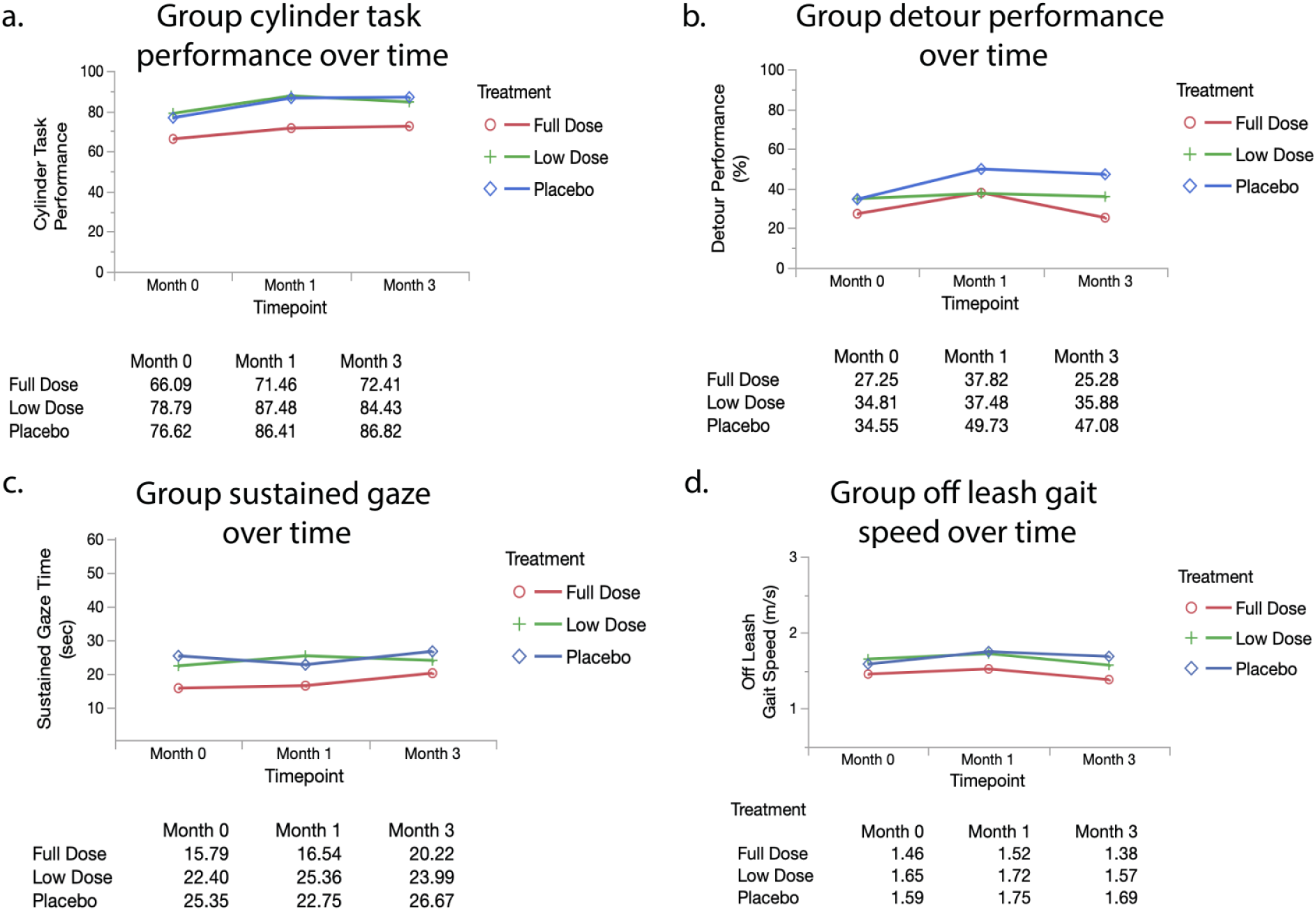
Repeated measures analyses of secondary outcome measures (Cylinder Task, Detour, Sustained Gaze, Off Leash Gait Speed). Mean values (adjusted as necessary) are reported at each timepoint by group. Group scores over time were compared in a repeated measures model (MANOVA analysis in JMP). The Wilks’ lamda value was evaluated with p<0.05 indicating a significant difference between groups. All original outcome measure data is provided in Supplementary Table S5.

#### Detour

Detour is a further challenge to the cylinder task, requiring flexibility in problem solving, which becomes more challenging as dogs age [56]. Dogs had lower baseline scores on detour as compared to the cylinder task. Similar to the cylinder task, the full dose group started out with a lower baseline score (27.3%) compared to both the low dose (34.8%) and placebo group (34.6%); these scores again were not significantly different (p=0.52) (Table 1). In the repeated measures analysis of change in score over time, detour required correction for fractional lifespan. The groups did not differ significantly across the 3 months (p=0.85) (Figure 6b). We saw a small decrease in score (decline in performance) with the full dose group at the primary endpoint.

#### Sustained Gaze

Sustained gaze is a test of focus and attention, with aging dogs shown to decline in the time in which they can maintain gaze for a treat [21]. At enrollment, there was no significant difference in performance between groups (p=0.22) (Table 1). No covariates reached significance to be included in the repeated measures analysis. While sustained gaze duration increased in all three groups at the primary endpoint, indicating an improvement, there was no significant difference between groups over time (p=0.59) (Figure 6c).

#### Off Leash Gait Speed

Off leash gait speed is both a measure of physical ability and motivation (for a treat). This has been shown to decline with age in both dogs and humans [30]. There was no significant difference in off leash gait speeds across groups at enrollment (p=0.55) (Table 1). Gait speed did not require correction for any covariates in the repeated measures analysis. All groups showed little change in speed, with no significant difference between treatment groups over time (p=0.59) (Figure 6d).

### Secondary Endpoint

Similar to the primary outcome measures, we assessed whether individuals were able to maintain their level of frailty, cognitive testing scores and gait speed from months three to six by comparing change in score across groups. We found the majority of dogs remained static (median change of approximately zero) across all groups from month three to month six (Supplementary Table S2). There was no significant difference between groups for frailty (p=0.80), inhibitory control (p=0.33), detour (p=0.28), sustained gaze (p=0.33) or off-leash gait speed (p=0.64).

#### Owner assessed happiness

Owners were asked to categorize their dogs’ level of happiness as the same, improved or deteriorated compared to the previous visit at three and six months. At three months, 10/18 (56%) owners with dogs in the full dose group reported static happiness, 8/18 (44%) improved and none reported a deterioration (Supplementary Table S5). While not significant (p=0.34), this differs from the low dose and placebo groups in which 2/21 (10%) and 3/20 (15%) reported a deterioration. At six months, 6/17 (35%) owners with dogs in the full dose group and 8/17 (47%) in the low dose group reported an increased level of happiness. Whereas only 4/17 (24%) of owners in the placebo group reported an increase. These differences were not significant (p=0.42).

#### Adverse Events

All dogs who received supplements/placebo were included in the adverse events analysis. At each visit, owners completed a checklist regarding possible adverse reactions (Supplementary Figure S6). There were 245 adverse events reported, ranging in VCOG grade: 1 (n=190), 2 (n=40), 3 (n=7), 4 (n=1), 5 (n=6). Events were only included if they were newly observable (not present at baseline) during the course of treatment. Of these reported events, 76 were in the placebo group, 102 in the low dose group and 67 in the full dose group (Supplementary Table S7). These events were evenly distributed across the groups.

Most events observed were VCOG grade 1 (n=190) or 2 (n=40). Grade 1 events were considered minimally disruptive by the owner, required no intervention, and resolved quickly on their own. Grade 2 events moderately impacted the patient’s daily life and usually necessitated outpatient veterinary care (antibiotics, pain management, minimally invasive procedure). There were seven severe (Grade 3) events that required medical intervention and had a significant effect on daily activities but were not immediately life threatening. These events were reported to us and medically managed by each patient’s primary care veterinarian. None of these events could be related to LYD6/2. Only one event was graded as four (requiring urgent medical attention): this dog presented to their final visit with lethargy, weakness and ascites. Over the course of the trial, six dogs developed disease that ultimately led to euthanasia or death (Grade 5); two in the placebo group, three in the low dose group and one in the full dose group. The reasons for these deaths included: severe respiratory disease, neoplasia, poor quality of life (severe cognitive dysfunction and incontinence) and surgical complications during a gastric dilation volvulus procedure.

Adverse events occurred across a variety of systems for all groups (Supplementary Table S7 and Figure 7). Only two adverse events occurred in the treatment groups that were not also seen in the placebo group (hypertension and an anaphylactic reaction). The hypertension was pre-existing and well controlled prior to the trial, but over the course of the trial, the dog developed worsening hypertension and was eventually euthanized due to poor quality of life. The anaphylactic reaction was in response to an antibiotic prescribed for a UTI which quickly resolved after discontinuation of the antibiotic. Therefore, it is unlikely that either of these events were a direct result of LY-D6/2.

**Figure 7:**
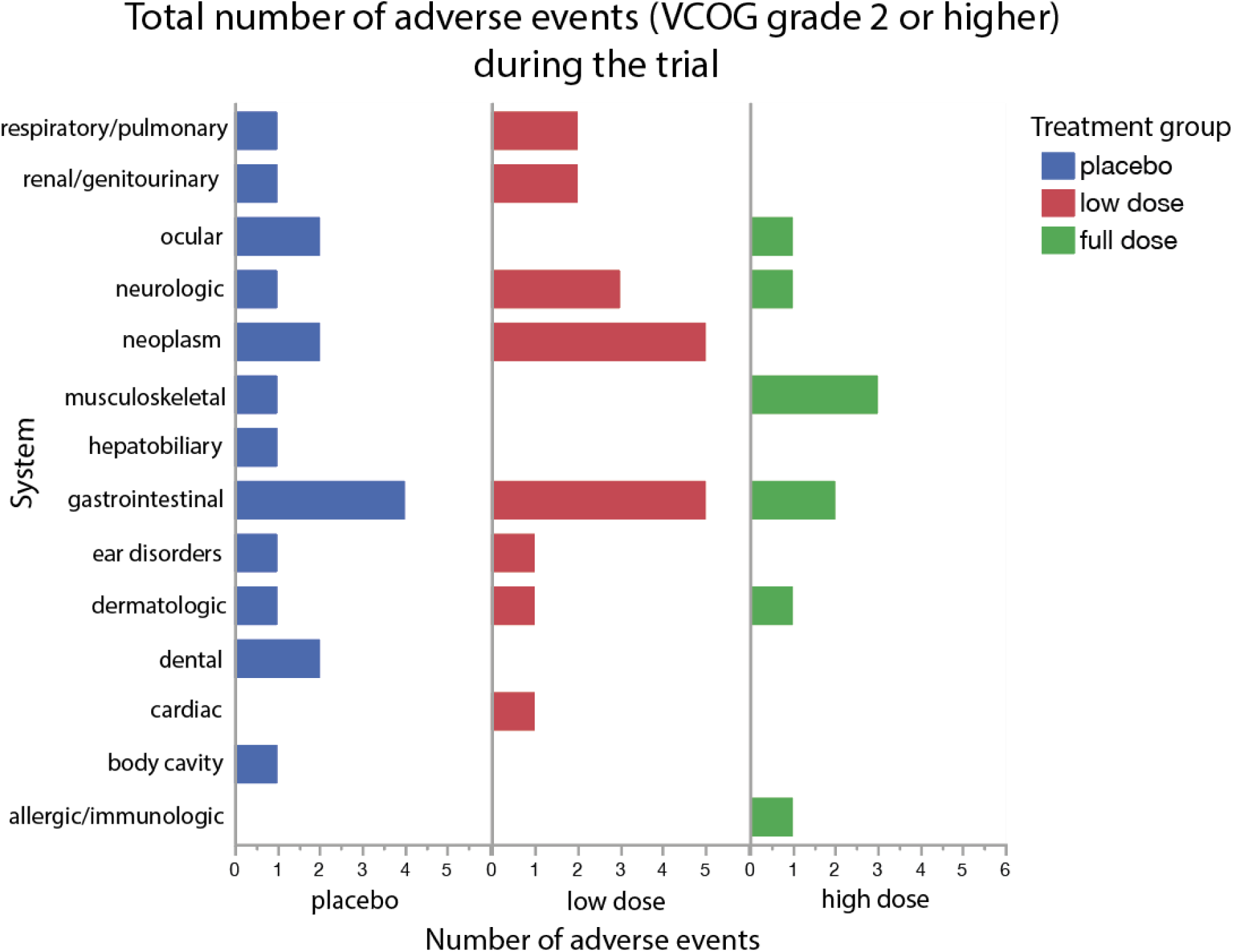
Total number of adverse events (VCOG grade 2 or higher) during the trial by group. Events were reported by the owner to be new during the trial (not present during/before baseline) and separated by system. Events were summed based on the system affected. All original adverse event data is provided in Supplementary Table S7.

There were 284 lab work changes over the course of the trial (Supplementary Table S7) and of these, only two were graded as VCOG 2 and one was graded as VCOG 3. The rest were mild, mostly incidental findings. The most common findings were anemia (n=21), lymphopenia (n=21) and hematuria (n=26). Any observed abnormalities were reported to the owner and pursued by the primary care veterinarian at their own discretion. Each group experienced a similar number of lab work abnormalities; placebo: 95, low dose group:106 and full dose group:87.

## Discussion

This randomized controlled blinded trial is one of the first of its kind, evaluating an anti-aging supplement that targets two hallmarks of aging in senior dogs. This clinical trial used a pragmatic approach that included dogs with mild to moderate cognitive impairment who met age, weight and relatively broad health criteria. Dogs were followed to a primary endpoint at three months, representing nearly two human years, and a secondary endpoint at six months, representing approximately three and a half human years. There was a significant difference in change in CCDR score across treatment groups from baseline to the primary endpoint at three months, with the largest decrease in score in the group of dogs who received the full dose of LY-D6/2. There was no difference between groups in changes in activity level, gait speed or in-house cognitive testing. However, while not significant, a higher percentage of dogs in the placebo group showed a deterioration in frailty (45.0%) compared to both the low and full dose LY-D6/2 groups (23.8% and 27.8%, respectively). The LY-D6/2 supplements were well tolerated; while many health issues occurred in this group of old dogs over the six-month trial period, there was no difference in prevalence of adverse events between treatment groups and none could be attributed to LY-D6/2.

The rationale of targeting two molecular mechanisms of aging was both to enhance the intervention strategy and to broaden the population of dogs that might benefit from the supplements. Both cellular senescence and depletion of cellular NAD+ are well established molecular hallmarks of aging and manipulation of both events in either direction has been shown to either ameliorate or worsen the aging phenotype in an experimental setting [37]. The use of a variety of different NAD+ precursors to increase cellular NAD+ concentrations is one of the most popular anti-aging strategies available at this time [53]. While there are now multiple studies demonstrating the ability to increase blood cell concentrations of NAD+ with oral supplements, and some evidence of a health benefit in middle aged and healthy old people [58,59,60], definitive evidence of an effect on the aging phenotype in people is still lacking [53] and the only data in dogs relates to a model of Duchenne’s muscular dystrophy [54]. Cellular NAD+ attrition interfaces with other mechanisms of aging and the possibility of a synergistic effect of NAD+ precursors when combined with another therapy has been proposed [55]. There has been particular interest in the rather complex interplay between cellular senescence and NAD+ depletion. It is clear that the SASP production by senescent cells is both metabolically taxing and reduces NAD+ concentrations by increasing CD38 expression [61]. In this clinical trial we chose to maximize the potential benefit of the anti-aging intervention by combining a senolytic with an NAD+ precursor.

Given the dearth of published placebo controlled randomized clinical trials examining anti-aging supplements in aged companion dogs, there is insufficient data available on parameters such as anticipated death rates, frequency and impact of comorbidity development, and size of caregiver placebo effect. We powered this clinical trial around detection of change in owner-quantified change in cognitive function. We used preliminary longitudinal data on dogs with mild to moderate cognitive impairment and made the observation that the vast majority of such dogs decline over a six-month period. This is supported by other published studies [62] and prevention of this deterioration would represent a meaningful benefit to dogs and their owners. Thus, the clinical trial was designed to detect a 50% reduction in the proportion of dogs that would show a deterioration in owner reported cognitive status. A primary endpoint at three months was chosen because of concerns about case attrition due to development of conditions such as cancer in this elderly population of dogs.

Clinical trials can be explanatory or pragmatic, with explanatory trials answering the question of whether an intervention is effective in a very specific patient population and pragmatic trials determining whether a therapy will be effective under normal conditions [63,64]. In this clinical trial, the study population was intentionally wide and the primary outcomes (owner reported cognitive status and activity detected by a collar mounted PAM) were straightforward to obtain and were directly relevant to a dog’s day-to-day life. The inclusion of simple owner reported assessments of improvement or deterioration in activity levels and happiness allowed identification of meaningful changes by owners blinded to treatment group. The results of this pragmatic clinical trial can immediately inform recommendations by primary care veterinarians.

In dogs, cognitive decline is strongly age-associated, with prevalence estimates of mild cognitive impairment in 28% of 11-12 year olds, and up to 68% of 16 year olds [65,66]. The odds of developing canine CCDS increase by 52% with each additional year of age [67]. There are several different validated owner questionnaires to quantify canine cognitive decline and the development of CCDS. The initial power analysis and patient recruitment was performed using CADES because this scale identifies mild cognitive changes through an option to choose an event frequency of q6 months. However, owner scores of cognition were solicited at baseline, one, three and six months and so the CCDR was chosen as a more reliable means of sampling cognitive status repeatedly within six months. Using this measure, only 2/18 dogs (11.1%) in the full dose group showed a deterioration in score compared with 6/21 (28.6%) in the low dose group and 8/20 (40%) in the placebo group. Moreover, the repeated measures analysis, using a model that accounted for frailty status at study start, showed a significant effect of supplementation with the full dose group showing the largest decline in score (clinical improvement). This effect was not maintained through six months with CCDR scores plateauing or increasing slightly in all three groups. Given this time period represents nearly 3.5 years of human life, this is perhaps not surprising.

Owners of 60% of dogs in the placebo group documented an improvement in cognition over a three-month period suggesting that there was a sizable and long-lasting caregiver placebo effect [68]. This caregiver placebo effect could be the result of optimism on the part of the owner; in addition, many owners participating in a longitudinal study of aging with our research group report that the interactions their dogs have when they visit the research site improve their attitude and engagement. Thus, this trial might be capturing the effect of increased social interaction and problem solving in elderly dogs through study participation. This is supported by studies in purpose bred dogs in which behavioral enrichment was associated with preservation of learning [69,70], through elevation of brain levels of BDNF expression [71].

The results of the secondary cognitive tests of attention and executive control performed at the research site did not differ between groups, with most groups showing a slight improvement. These cognitive tests have not been used in a longitudinal clinical trial previously, and it is possible that these dogs learned how to perform the tests better with each visit. In addition, we might again be capturing the positive effect of trial participation.

Assessing activity levels in senior and geriatric dogs is challenging because they tend to lead quite sedentary lives and, as for all companion dogs, their activity is strongly influenced by their owner’s activity [72,73]. We used collar mounted activity monitors to collect data over a two-week period at each evaluation point, and evaluated weekdays and weekends separately to allow for owner schedules differing over the weekend [74]. The baseline data showed the typical peaks in activity when the household rises in the morning and comes home from a day of work during the week. We first performed an FLM analysis as this captures and smooths data allowing a per-minute comparison between groups over a 24-hour period. Given that multiple comparisons are made when performing an analysis in this way, only group differences that reach the threshold of global significance are compelling, while those that reach pointwise significance highlight areas of interest. However, this analysis did not reveal any pattern of consistent change between groups. It was interesting to note that morning and evening activity on weekdays did increase in all three groups, again suggesting either placebo or a true beneficial effect of trial participation. Given these elderly dogs are not very active, but were reported to demonstrate small bursts of energy by owners, we also compared the sum of the total daytime and nighttime activity counts for each dog between groups. Unlike the FLM, which cannot be performed with covariates, this allowed us to build a model taking covariates into account, but no differences were noted between groups. Finally, we compared the owner reports of activity changes and more full dose group owners noted an improvement than the other groups. This raises the question of how best to capture activity changes in elderly dogs.

Frailty is a well described syndrome in aging people but descriptions of frailty are few in dogs [75,76,77]. We have developed a rapid screening tool for frailty that combines owner responses to questions around the five key frailty domains, and assessment of body and muscle condition score by the research group [78]. While there was no significant difference between groups in percentage of frail individuals at study start, approximately 35% of the placebo group were frail compared with 48% in both treatment groups. One would anticipate that aged dogs would become frailer over time, but in our study, the majority of dogs in all three groups improved in their frailty status over the first three months with a lower percentage of dogs in the treatment groups deteriorating than in the placebo group. Currently there is a dearth of data following frailty in elderly dogs longitudinally and so it is difficult to comment on whether this is unusual, but once again, it does suggest either a placebo effect, or a positive effect of trial participation. This study has some weaknesses, first, this was a pragmatic clinical trial with no attempt to target dogs who exhibited higher levels of senescence or lower levels of NAD; a more targeted trial might be better able to capture changes due to alterations in these systems. Further, biomarkers of senescence and NAD+ levels were not measured. Future studies may benefit from incorporating these biological markers as outcomes. Our inclusion of cognitive tests was designed to capture changes in performance measures (as compared to owner-rated changes in signs) however these tests have not been used longitudinally, and while correlations have been seen between some of these measures and owner-questionnaires [79], their responsiveness as outcome measures was unknown. Dogs may learn to perform these tasks with practice, and additional longitudinal data will be needed to understand how performance on these tasks changes with time. Finally, the full dose group was significantly older than the other groups, although fractional lifespan did not differ. Cognitive decline is associated with age rather than fractional lifespan, and so this could have influenced the outcomes negatively for this group. However, age was examined in the univariate analysis of CCDR change and was not associated with the outcome.

We conclude that LY-D6/2 can be used safely to mitigate cognitive decline in senior dogs and might have broader effects on dog health manifesting as improved happiness and reduced frailty. This trial highlights the viability of targeting hallmarks of aging to impact the health of aging dogs. The pragmatic design means the results are immediately applicable to an aging dog population and are potentially relevant to people. Clinical trial participation appears to be beneficial in an aging dog population.

## Methods

This blinded, randomized, controlled (RCT) clinical trial was conducted and reported according to the CONSORT and ARRIVE Guidelines with the approval of the North Carolina State University Institutional Animal Care and Use Committee under protocol # 21-376-O. All procedures were performed in accordance with these approved protocols and institutional guidelines. Owners of the dogs who participated in these studies reviewed and signed an informed consent form. IRB approval was not sought because all collected data pertained to dogs and as such the work was categorized as “Non-Human Subject Research”.

### Study design

This 3-arm RCT was designed to evaluate the effect of LY-D6/2^TM^ on cognitive function and activity in aged dogs over a six-month period. The primary endpoint of the study was three months (representing approximately 1.75 human years) and the secondary endpoint was six months (representing approximately 3.5 human years). The primary outcomes were change in owner assessed cognitive function (via Canine Cognitive Dysfunction Rating (CCDR) scale) [80] and activity level (via a collar mounted physical activity monitor (PAM)) [81]. Secondary outcomes included changes in frailty phenotype, attention (sustained gaze), inhibitory control (cylinder task), cognitive flexibility (detour task) and off-leash gait speed. Preliminary data from six mild to moderately cognitively affected dogs (based on owner assessment using the Canine Dementia Scale (CADES) [62] were used to perform the power analysis. Five of these six dogs showed a deterioration in owner reported cognitive function over a six-month period. It was determined that 20 dogs per group would detect a 50% reduction in the number of dogs who show cognitive deterioration with a power of 80%. Ten additional dogs were included in order to account for attrition. Dogs were randomized in blocks of 9 (3 per group) by the NC State pharmacy using a random number generator. Investigators and owners were blinded to treatment identity until data analysis was complete. The timeline of participation is provided in Supplementary Figure S8.

### Inclusion and exclusion criteria

In order to participate, dogs had to be greater than or equal to 10 years of age, weigh more than 8kg, and be able to walk independently with sufficient hearing and vision to be able to perform cognitive tests. They had to be treat-motivated, willing to engage with the investigators, and show signs of mild to moderate cognitive impairment (based on CADES score). Owners had to be able to complete online questionnaires, administer the supplements, keep an activity monitor on their dogs for two-week periods and attend scheduled appointments.

Dogs with comorbidities likely to significantly adversely affect health over the course of the clinical trial were excluded. Examples include metastatic neoplasia, hyperadrenocorticism, diabetes mellitus, congestive heart failure, and refractory epilepsy. Aggressive dogs were excluded as they could not safely perform the cognitive tests. Dogs with evidence of an active urinary tract infection (clinical signs, bacteriuria and pyuria) were excluded until treated.

### Intervention

The investigational veterinary product (IVP), LY-D6/2 was a proprietary combination of a senolytic and an NAD+ precursor. Dogs were administered placebo, low or full dose LY-D6/2 starting the day following Day 0 assessment. The doses were determined in pharmacokinetic and safety studies performed by Animal Bioscience. Owners were instructed to administer the NAD+ precursor (or placebo) capsule(s) once daily and the senolytic (or placebo) capsule(s) on two consecutive days each month. Owners were given customized calendars to record administration of the capsules throughout the study. Both these calendars and capsule counts (checked at each visit) were utilized to determine compliance.

### Recruitment and Eligibility

Dogs were recruited from the local community through postings on the NC State CVM clinical trials website and social media from January 2022 through June 2023. Dogs were initially screened via an online survey platform (Qualtrics) and those meeting age (≥ 10 years), weight (≥ 8 kgs) and cognitive status (CADES category mild or moderate) requirements proceeded to a telephone interview to confirm cognitive changes were age-related and that the owner understood the clinical trial design and commitments. Eligible participants proceeded to an in-person screening at the NC State College of Veterinary Medicine. Owners reviewed and signed informed consent forms at this time. The screening visit included physical examination, lab work (CBC and serum biochemistry panel) and behavioral assessment to confirm eligibility prior to enrollment. A PAM (WGT3X-BT monitor, Actigraph, Pensacola, FL) was placed on the dog’s collar and the owners were provided with a log to record times where the collar/PAM was removed and/or changes from routine activity. The appointment for the first day of the trial was scheduled two weeks after the screening appointment in order to allow for adequate PAM data collection prior to beginning treatment (Supplementary Figure S8). If a UTI was detected at screening, dogs were treated with an appropriate antibiotic by their primary veterinarian and had to have a clear urinalysis prior to entering the clinical trial.

### Data collection

Study data were collected and managed using REDCap electronic data capture tools hosted at NC State University [82,83].

Questionnaires were distributed to owners via REDCAP (https://www.project-redcap.org/) seven days prior to each appointment. Basic signalment and medical history were obtained. Owners were also instructed to indicate any changes in pet health, medication, environment and attitude. RedCap questionnaires included CCDR and frailty phenotype. The CCDR Questionnaire consists of 13 behavioral items related to orientation, memory, apathy, olfaction and locomotion. (Supplementary Figure S6). Scores from each item are summed (16-80), where a score of 50 or greater indicates CCDS[80]. The frailty phenotype screening tool assesses age-related deficits in mobility, exhaustion, social activity, muscle condition, and nutritional status and predicts six-month mortality[78]. Criteria have been established to classify an individual as impaired in each of these five domains and the individual was classified as frail if criteria are met for three or more of these domains. (Supplementary Figure S6) These data were expressed both as frail, yes or no and as scores from 0-5 (number of impaired domains). In addition to chronological age, fractional lifespan was also calculated for each dog using an equation that incorporates dog height and weight [84]. In order to evaluate the owner’s perception of efficacy, each was asked to categorize their dog’s mobility and their level of happiness as improved, the same or worse at months three and six.

Dogs underwent a physical, neurological and orthopedic examination (Supplementary Figure S6). Joint and spinal pain was quantified using an established scale, where scores are summated to give a score from 0-76 [85,86]. Off leash gait speed over a five-meter distance was measured in triplicate and the mean value was calculated [30]. Cognitive testing included the sustained gaze task and two tests of executive function (cylinder and detour tasks). In the sustained gaze task, dogs are asked to focus on a treat held by the handler’s face and the time they maintain the gaze is recorded, performed in triplicate for calculation of the mean value in seconds [21]. The cylinder and detour tasks both generate outcomes as a percentage of correct choices in a set of trials (n=8). All tasks are described in detail elsewhere [79]. All findings from examinations, mobility tests and cognitive tests were recorded in the REDCap database.

These assessments were performed at time 0, and at one, three and six months. Blood work and urinalysis was repeated at one and six months, and a urine dipstick test was performed at three months. All lab work performed during the study are provided in Supplementary Table S9. The PAM device was placed for two weeks at the screening appointment and again at months one, three and six. It was removed after two weeks and returned to the research group along with the owner log that recorded collar removal or unusual activity. Data were downloaded for analysis after each time period using ActiLife software (version 6.13.3; Actigraph, Pensacola, FL) All raw activity data are provided in Supplementary Table S10.

Changes in the dog’s environment (e.g. address change, addition of another pet or other household changes) were categorized as present or absent at one, three and six months as were changes in medications the dogs were receiving. Adverse events (changes in blood work and/or new clinical signs) were categorized as grade 1 through 5 according to the VCOG-CTCAEv2 guidelines [87]. Those classified as grade 2 or above were reviewed by the safety monitor and the treatment mask broken if considered appropriate.

### Statistical analysis

All statistical analyses were performed with patient and group identity masked. Analyses were performed using JMP Pro 16 (SAS Institute, Cary, NC). Summary statistics were generated for dog demographics in each group at time of study entry. Continuous data were reported as mean and standard deviation (SD) or median and range depending on data distribution; categorical and ordinal data were reported as population proportion. Data were examined for normality based on inspection on Q-Q plots, histogram and Shapiro-Wilks test. Baseline demographic differences between groups were assessed using one-way ANOVA or Wilcoxon Rank Sum test (based on data normality) for continuous data. For ordinal/categorical data (Sex, BCS, Frailty Status [Y/N], Frailty Score) contingency tables were constructed and a chi-square test was performed with Fisher Exact test performed if needed. Changes in household and changes in medication were assessed in a similar manner, utilizing a chi-square test to assess significance (p<0.05).

Changes in our owner-based assessments (CCDR and frailty phenotype) from time zero to the primary endpoint (month three) were categorized as success (unchanged or decreased score) or failure (increased score). These categorical changes in CCDR and frailty were compared between groups by constructing contingency tables and performing a chi-square test. I.

Changes in CCDR score, cognitive testing performance and off-leash gait speed over the study were compared between groups with repeated measures ANOVA. Repeated measures ANOVA requires approximately normal distribution, so the residuals of these data within the model were assessed for distribution and outliers based on inspection of Q-Q plots and histogram. To account for covariates that might influence outcomes in senior dogs, univariate analyses of age, fractional lifespan, weight, sex (M or F), BCS, baseline frailty status (frail in 3 domains, yes or no), change in household and change in medication was performed for each of the outcome measures. Those covariates that reached significance of p<0.1 were incorporated into the repeated measures ANOVA model for that outcome in addition to treatment effect. Overall differences between groups over time were assessed via wilk’s lambda using a threshold of p<0.05.

In order to assess whether any treatment effect was maintained from month three to month six, changes in individual CCDR, Frailty, cognitive testing and off leash gait speed were calculated (month 6-month 3) and compared across groups. Changes were assessed via a Wilcoxon test, with p<0.05 indicating significance. Owner’s perception of mobility and happiness level (categorized as improved, static or worse) were evaluated at months three and six via a chi-square test

Activity data for weekdays and weekends were analyzed separately given the impact of owner schedule on dogs’ activity [16,74]. Data underwent quality control to identify the first five weekdays and two weekend days with complete data. Data from days that owners reported removal of the collar or unusual activity and days that included more than three consecutive hours with no (zero) activity detected were excluded. The mean activity per minute was calculated from five weekdays and from two weekend days and the change in mean activity/minute was calculated for each individual dog between study start and the primary end point at month three. The change between three and six months was also calculated. Baseline and change in activity were analyzed using functional linear modeling (FLM) in order to facilitate modeling of activity data over time, thus avoiding the data loss that occurs when periods of time are averaged [16,81]. This was performed using RStudio 2021 (PBC, Boston, Ma) package “Actigraphy” version 1.4.0, which uses a Fourier expansion model to transform raw data into smoothed activity curves. Differences between groups were evaluated using a non-parametric permutation F-test with 1000 permutations performed. This generates a curve of the F-statistic over time. Point-wise (a curve with the F permutation proportion at each time point) and global (single number referring to the proportion of maximized F values from each permutation) critical F values are calculated to set thresholds for statistical significance [88].

The FLM analysis only allows the incorporation of a single covariate (treatment group) and therefore cannot account for other potential confounding influences that we predict exist in the senior pet population. For this reason, we also analyzed total day time (5:00am-10:59pm) and night time (11:00pm-4:59am) activity in a repeated measures analysis (similar to the rest of the outcome measures). Covariates (age, fractional lifespan, weight, sex, BCS, frailty status, change in household and change in medication) were incorporated if they reached significance (p<0.1) in a univariate analysis. As with the FLM analysis, weekday and weekend data were assessed separately and wilks lambda values were assessed for significance (p<0.05). Change in cumulative activity during days and nights on weekdays and weekends across groups was assessed via a Wilcoxon test with p<0.05 representing significance.

## Acknowledgements

This clinical trial was funded by Animal Bioscience, Boston MA. Animal Bioscience played no role in data acquisition, analysis or presentation.

We would like to acknowledge all the dogs and their owners and Amanda Valentino and Katerina Slaughter for assistance with dog handling.

## Author Contributions

KES, NJO, MEG and KR were responsible for the design, analysis, and primary writing of the manuscript for this study. KES, AM, KR, BC, CW, ZA, CY all participated in the data acquisition; all authors participated in editing and review of the manuscript.

## Data Availability Statement

All data analyzed in this study has been provided in the supplementary data as Supplementary Data files Supplementary Tables: S5, S9 and S10.

---

Supplementary Figure S3: Repeated measures analysis of summated activity monitor activity by group. Repeated measures analysis of summated activity across month 0, 1 and 3. Day (5am-10:59pm) and night (11pm-4:59am) were assessed separately, as were weekdays and weekends. Mean summated activity levels by group were obtained from repeated measures models (MANOVA analysis in JMP) with adjustment for covariates when necessary. The Wilks’ lamda value was evaluated, with p<0.05 indicating a significant difference between groups. All original activity data is provided in Supplementary Table S10.

Supplementary Figure S6: Study Exam Forms and Questionnaires. Forms are displayed in the following order: physical examination form, neurological examination form, orthopedic examination form, Canine Cognitive Dysfunction Rating (CCDR) Questionnaire, Frailty Assessment, Changes since last visit.

Supplementary Figure S8: Study Timeline. The timeline of all visits in the study are provided along with all assessments performed at each visit. Activity monitors were placed at specific visits in order to obtain two week increments of activity data for each timepoint. Intervention was started the day following an individual’s baseline (month zero) visit.

