## Supplementary material for "A Randomized, Controlled Clinical Trial Demonstrates Improved Cognitive Function in Senior Dogs Supplemented with a Senolytic and NAD+ Precursor Combination": Supplementary Files.pdf

| | Time Period | Placebo (n= 20, 18) | Low Dose (n= 21, 17) | Full Dose (n= 18, 18 ) | p value | Fisher's Exact Test |
|---|---|---|---|---|---|---|
| Changes to Household | Month 0 to Month 3 | Yes: 7<br>No: 13 | Yes: 2<br>No: 19 | Yes: 4<br>No: 14 | 0.13 | 0.15 |
| | Month 3 to Month 6 | Yes: 3<br>No: 15 | Yes: 1<br>No: 16 | Yes: 4<br>No: 14 | 0.35 | 0.50 |
| Changes to Medication | Month 0 to Month 3 | Yes: 6<br>No: 14 | Yes: 8<br>No: 13 | Yes: 3<br>No: 15 | 0.24 | |
| | Month 3 to Month 6 | Yes: 4<br>No: 15 | Yes: 4<br>No: 14 | Yes: 3<br>No: 17 | 0.32 | |

Supplementary Table S1: Household and medication changes in participants over the course of the study by group.

| | Placebo (n=17) | Low Dose (n=17) | Full Dose (n=17) | p value |
|---|---|---|---|---|
| CCDR Score | 0<br>(-11 - 7) | 0<br>(-11 - 14) | 1<br>(-6 - 26) | 0.44 |
| Frailty Score | 0<br>(-2 - 2) | 0<br>(-1 - 2) | 0<br>(-1 - 3) | 0.80 |
| Cylinder Task (Inhibitory Control) (%) | 0<br>(-25 - 25) | 0<br>(-50 - 37.5) | 0<br>(-75 - 37.5) | 0.33 |
| Detour (%) | 0<br>(-75 - 37.5) | 0<br>(-87.5 - 37.5) | 0<br>(-25 - 37.5) | 0.28 |
| Sustained Gaze (sec) | -4.24<br>(-42.53 - 22.37) | 2.18<br>(-25.82 - 17.34) | -1.11<br>(-20.25 - 29.28) | 0.33 |
| Off-Leash Gait Speed (m/s) | 0.02<br>(-0.83 - 0.99) | -0.08<br>(-0.68 - 0.78) | 0.05<br>(-0.66 - 0.43) | 0.64 |

Supplementary Table S2: Median (Range) change in outcome measures (Month 6 - Month 3) by group.

a.

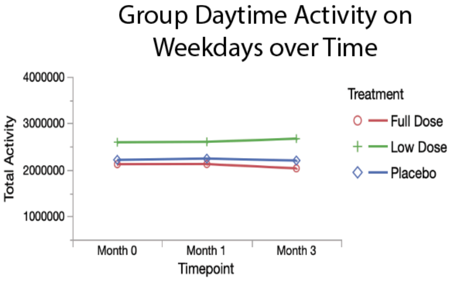

| | Month 0 | Month 1 | Month 3 |
|---|---|---|---|
| Full Dose | 2121177.33 | 2124223.24 | 2032127.19 |
| Low Dose | 2594228.34 | 2602078.25 | 2669909.66 |
| Placebo | 2215007.51 | 2241396.15 | 2200677.55 |

b.

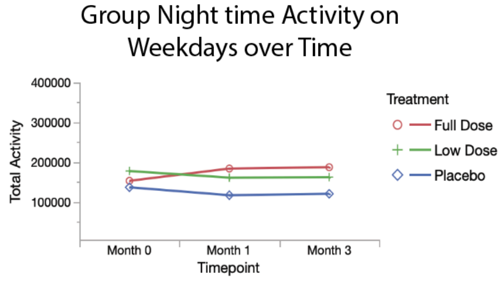

| | Month 0 | Month 1 | Month 3 |
|---|---|---|---|
| Full Dose | 152509.50 | 183458.48 | 186752.90 |
| Low Dose | 177181.46 | 160498.97 | 161784.77 |
| Placebo | 136192.49 | 116236.34 | 119815.79 |

c.

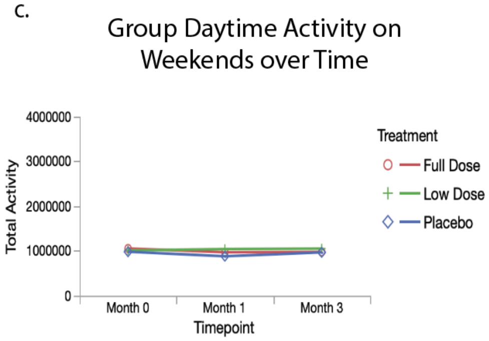

| | Month 0 | Month 1 | Month 3 |
|---|---|---|---|
| Full Dose | 1056012.75 | 971746.05 | 979355.00 |
| Low Dose | 1012168.03 | 1042947.72 | 1053542.23 |
| Placebo | 982300.10 | 881980.71 | 968383.09 |

d.

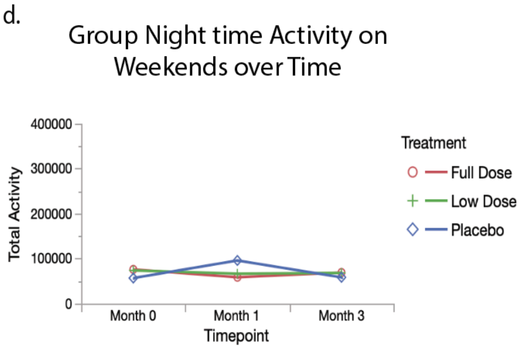

| | Month 0 | Month 1 | Month 3 |
|---|---|---|---|
| Full Dose | 75983.81 | 58541.70 | 69125.24 |
| Low Dose | 73766.51 | 66468.32 | 68212.73 |
| Placebo | 56743.98 | 95902.81 | 58369.47 |

Supplementary Figure S3: Repeated measures analysis of summated activity monitor activity by group.

Repeated measures analysis of summated activity across month 0, 1 and 3. Day (5am-10:59pm) and night (11pm-4:59am) were assessed separately, as were weekdays and weekends. Mean summated activity levels by group were obtained from repeated measures models (MANOVA analysis in JMP) with adjustment for covariates when necessary. The Wilks' lamda value was evaluated, with $p<0.05$ indicating a significant difference between groups. All original activity data is provided in Supplementary Table S10.

| | | | Cumulative Change in Activity | p value |
|---|---|---|---|---|
| Weekday Activity | Daytime | Placebo (n=16) | 124158 (-954337 - 632044) | 0.89 |
| | | Low Dose (n=16) | 142019 (-344941 - 1253340) | |
| | | Full Dose (n=17) | -25522 (-805479 - 753977) | |
| | Nighttime | Placebo (n=16) | 10092 (-25852 - 125521) | 0.10 |
| | | Low Dose (n=16) | 4609 (-98981 - 190055) | |
| | | Full Dose (n=17) | -17168 (-139014 - 102041) | |
| Weekend Activity | Daytime | Placebo (n=16) | 139652 (-589871 - 962594) | 0.60 |
| | | Low Dose (n=16) | 113584 (-395776 - 548630) | |
| | | Full Dose (n=17) | 7127 (-597520 - 398160) | |
| | Nighttime | Placebo (n=16) | -1871.64 (-75114 - 78950) | 0.83 |
| | | Low Dose (n=16) | 1822 (-65863 - 128574.97) | |
| | | Full Dose (n=17) | 7243 (-129861 - 167042) | |

Supplementary Table S4: Median (range) cumulative change in activity (month 6 - month 3) by group.

### The effect of combined supplementation on activity and attention in aging dogs

**Patient label**

**Study ID: _________ Date:_____________ Visit month #:___________ Estimated Life stage:_____________**

Weight (kg): __________ Temp: _________ Pulse: _________ Respiration: _________ MM/CRT: ________

Heart murmur (1-6): _________ BCS (1-9):_______

| | Normal | Abnormal | Describe |
|---|---|---|---|
| EENT/oral | | | |
| CVR | | | |
| GIT/ABD | | | |
| MSK | | | |
| LYMPH | | | |
| DERM | | | |
| GU | | | |
| Ortho | | | |

EXAMINATION NOTES:

______________________________________________________________________________

______________________________________________________________________________

UPDATES SINCE LAST VISIT

______________________________________________________________________________

______________________________________________________________________________

______________________________________________________________________________

### Neurologic exam

**Patient label**

Date:______________ Visit month #:_________

1. Mental Status:_______________ Alert Depressed Disoriented Stuporous Comatose Other
2. Posture:______________________ Normal Head Tilt Head Turn Other (describe)
3. Gait:_________________________ Normal Ataxia Paresis Plegia Lame Circling
4. Palpation: (0 = normal, 1 = mild, 2 = moderate, 3 = severe)

a.Neck pain: ____________ b.TL pain: _____________ c. LS pain: ___________ d. Head pain_________

5. Postural reactions: 0(absent), 1(reduced), 2(normal), 3(increased), 4(clonus)

| | LF | RF | LH | RH |
|---|---|---|---|---|
| CP | | | | |
| Hopping | | | | |

6. Cranial nerves: 0(absent), 1(reduced), 2(normal), 3(increased), 4(clonus)

| | L | R |
|---|---|---|
| Menace (II, VII) | | |
| Palpebral (V, VII) | | |
| Oculovestibular reflex (III, VI, VIII) | | |
| Gag reflex/Jaw tone (IX, X, V) | | |
| Eye Position/Strabismus (III – VL, VI – medial) | | |
| Facial muscles – eyelid, ear, lip, nose- (VII) | | |
| Masticatory muscles – (V) | | |
| Facial Sensation – (V) | | |
| Nystagmus – (VIII) | | |

7. Spinal reflexes: 0(absent), 1(reduced), 2(normal), 3(increased), 4(clonus)

| | LH | RH | LF | RF |
|---|---|---|---|---|
| Patellar reflex | | | | |
| Cranial Tibial reflex | | | | |
| Sciatic reflex | | | | |
| Flexor Withdrawal | | | | |
| Panniculus | | | | |
| Deep Pain | | | | |
| Perineal Reflex | | | | |

### Joint Evaluation

**Patient label**

Study ID:________ Date:_____________ Visit month #:________

| Forelimb | ROM | | Pain | | Crepitus | | Effusion | | Thickening | |
|---|---|---|---|---|---|---|---|---|---|---|
| | L | R | L | R | L | R | L | R | L | R |
| Manus | | | | | | | | | | |
| Carpus | | | | | | | | | | |
| Elbow | | | | | | | | | | |
| Shoulder | | | | | | | | | | |

| Hindlimb | ROM | | Pain | | Crepitus | | Effusion | | Thickening | |
|---|---|---|---|---|---|---|---|---|---|---|
| | L | R | L | R | L | R | L | R | L | R |
| Pes | | | | | | | | | | |
| Tarsus | | | | | | | | | | |
| Stifle | | | | | | | | | | |
| Hip | | | | | | | | | | |

Range of Motion:
**0**: normal; **1**: mild-moderate decreased; **2**: severely decreased

Pain based on manipulation:
**0**: Does not notice; **1**: Orients to site, does not resist or mild resistance (mild); **2**: Orients to site, slight objection to manipulation (moderate); **3**: Withdraws from manipulation, may vocalize, may turn to guard area (significant); **4**: Tries to escape/prevent manipulation, may bite or show aggression (severe)

Crepitus:
**0**: none, no crunching; **1**: mild, only feel crunching sometimes **2**: moderate, crunching felt always, may be painful;
**3**: severe, can feel and hear crunching, may be painful

Effusion:
**0**: none, no fluid pocket felt; **1**: mild, small fluid pocket felt only on palpation; **2**: moderate, prominent on palpation; **3**: severe, may see visible fluid pocket

Thickening:
**0**: none, can feel all anatomic structures easily; **1**: mild, less defined anatomic structures; **2**: moderate, can slightly define anatomic structures;
**3**: severe, can no longer feel anatomic structures

### Canine Cognitive Dysfunction Rating

Please indicate the frequency of your dog's behaviors.

| | Never | Once a month | Once a week | Once a day | > Once a day |
|---|---|---|---|---|---|
| 1. How often does your dog pace up and down, walk in circles and/or wander with no direction? | ○ | ○ | ○ | ○ | ○ |
| 2. How often does your dog stare blankly at the walls or floor? | ○ | ○ | ○ | ○ | ○ |
| 3. How often does your dog get stuck behind objects and is unable to get around them? | ○ | ○ | ○ | ○ | ○ |
| 4. How often does your dog fail to recognize people or pets? | ○ | ○ | ○ | ○ | ○ |
| 5. How often does your dog walk into walls or doors? | ○ | ○ | ○ | ○ | ○ |
| 6. How often does your dog walk away while, or avoid, being petted? | ○ | ○ | ○ | ○ | ○ |

| | Never | 1-30% times | 31-60% times | 61-99% times | Always |
|---|---|---|---|---|---|
| 7. How often does your dog have difficulty finding food dropped on the floor? | ○ | ○ | ○ | ○ | ○ |

**Please note the answer choices reflect the CHANGE in your dog's behavior over the past 6 months. If your dog has never exhibited the behavior, please select "the same".**

| | Much less | Slightly less | The same | Slightly more | Much more |
|---|---|---|---|---|---|
| 8. Compared with 6 months ago, does your dog now pace up and down, walk in circles and/or wander with no direction or purpose? | ○ | ○ | ○ | ○ | ○ |
| 9. Compared with 6 months ago, does your dog now stare blankly at the walls or floor? | ○ | ○ | ○ | ○ | ○ |
| 10. Compared with 6 months ago, does your dog urinate or defecate in an area it has previously kept clean? | ○ | ○ | ○ | ○ | ○ |
| 11. Compared with 6 months ago, does your dog have difficulty finding food dropped on the floor? | ○ | ○ | ○ | ○ | ○ |

| | | | | | |
|---|---|---|---|---|---|
| 12. Compared with 6 months ago, does your dog fail to recognize familiar people or pets? | ○ | ○ | ○ | ○ | ○ |
| | Much less | Slightly less | The same | Slightly more | Much more |
| 13. Compared with 6 months ago, is the amount of time your dog spends active? | ○ | ○ | ○ | ○ | ○ |

Total CCDR score ____________________

#### Frailty Phenotype Questionnaire and Scoring Guide for Population 2

---

#### 1. Nutrition:

- Body Condition Score (1 - 9)

  ▢ 1 ▢ 2 ▢ 3 ▢ 4 ▢ 5 ▢ 6 ▢ 7 ▢ 8 ▢ 9

- My dog now:

  ▢ Has an increased appetite compared to 6 months ago

  ▢ Eats normally (i.e. eats regular food, and finishes in the same amount of time)

  ▢ Will still eat their regular food but takes longer to finish

  ▢ Does not want to eat their regular food but will still eat treats or people food

  ▢ Acts interested in food but walks away from the bowl without eating

  ▢ Cannot be enticed to eat no matter what you offer

---

#### 2. Social Activity:

- Over the past week, how many days has your dog been playful?

  (No days) ▢ 0 ▢ 1 ▢ 2 ▢ 3 ▢ 4 ▢ 5 ▢ 6 ▢ 7 (Everyday)

---

#### 3. Exhaustion:

- How often does your dog rest (stop, sit, or lie down) during exercise?

  ▢ Never

  ▢ Hardly ever

  ▢ Occasionally

  ▢ Frequently

  ▢ Very frequently

---

#### 4. Mobility:

- On average, how frequently does your dog display weakness or lameness in his/her limbs when walking?

  *Weakness/ Lameness is defined as reduced strength in a limb. This can appear as limping (placing less weight on one leg compared to another), increased effort moving the limbs, scuffing of the paws, short strides, or falling.*

  - ▢ Not at all
  - ▢ Less than 10% of the time
  - ▢ 10 - 50% of the time
  - ▢ More than 50% of the time

- On average, how frequently does your dog display stiffness in his/her limbs when walking?

  *Stiffness is defined as reduced flexing of the joints. This can present as reduced bending of the limbs, shorter strides, or the dog appearing to be "straight-legged".*

  - ▢ Not at all
  - ▢ Less than 10% of the time
  - ▢ 10 - 50% of the time
  - ▢ More than 50% of the time

- Does your dog ever make mistakes while walking?

  *Mistakes may include scuffing of the paws, stumbling, swinging the limbs abnormally outward or inward such that they cross, a lack of coordination between the limbs, and/or skipping steps.*

  ▢ No ▢ Yes

#### 5. Muscle Condition:

- Forelimb Muscle Condition Score (0 - 3)

  ▢ 0 ▢ 1 ▢ 2 ▢ 3

- Hindlimb Muscle Condition Score (0 - 3)

  ▢ 0 ▢ 1 ▢ 2 ▢ 3

- Epaxial Muscle Condition Score (0 - 3)

  ▢ 0 ▢ 1 ▢ 2 ▢ 3

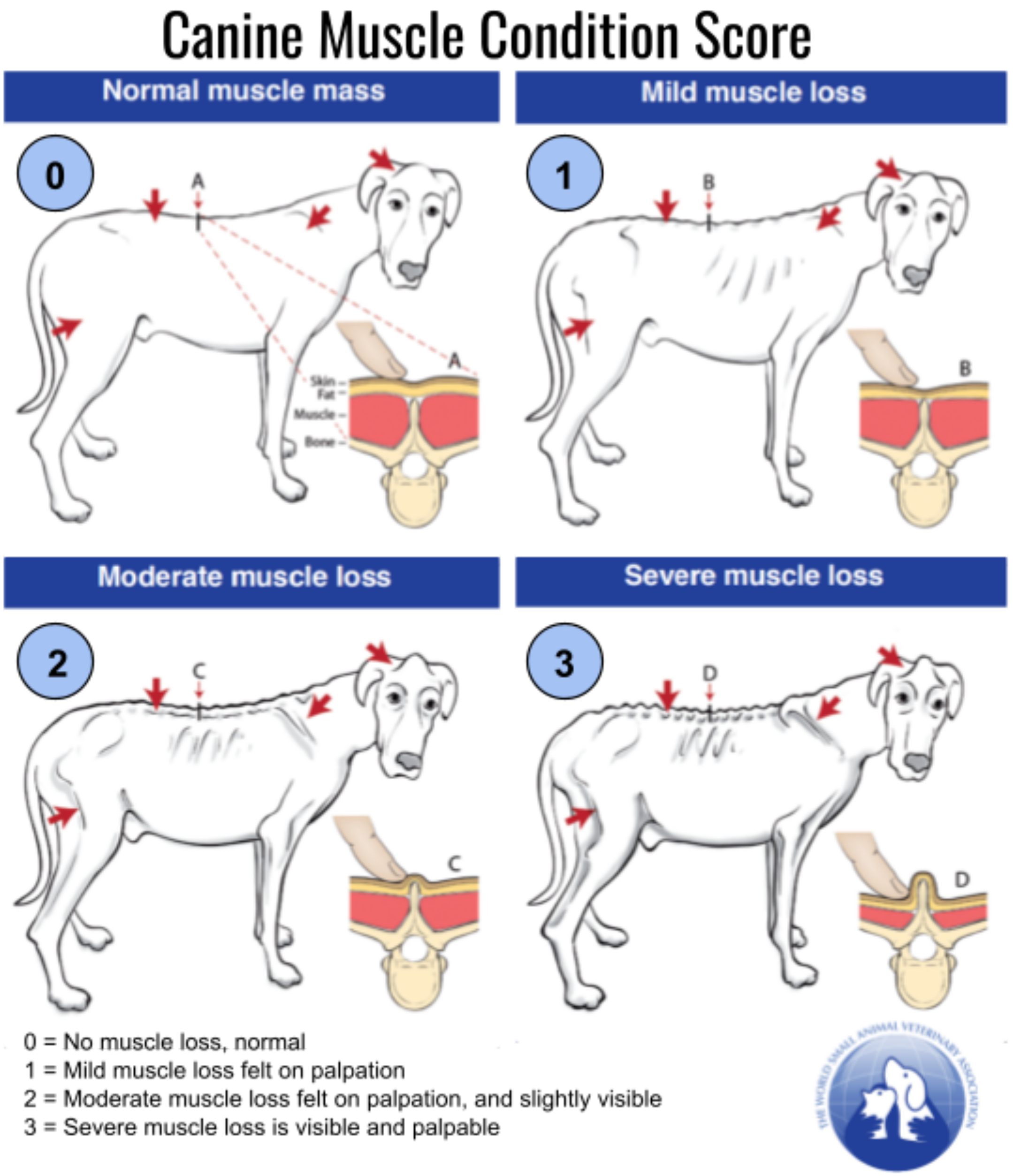

#### Frailty Phenotype Scoring

*Use the following key to complete the summary section and determine the total impaired domains*

---

##### 1. Nutrition:

- Body Condition Score (1 - 9)
  ▢ 1 ▢ 2 ▢ 3 ▢ 4 ▢ 5 ▢ 6 ▢ 7 ▢ 8 ▢ 9
- My dog now:
  - ▢ Has an increased appetite compared to 6 months ago
  - ▢ Eats normally (i.e. eats regular food, and finishes in the same amount of time)
  - ▢ Will still eat their regular food but takes longer to finish
  - ▢ Does not want to eat their regular food but will still eat treats or people food
  - ▢ Acts interested in food but walks away from the bowl without eating
  - ▢ Cannot be enticed to eat no matter what you offer

---

##### 2. Social Activity:

- Over the past week, how many days has your dog been playful?
  (No days) ▢ 0 ▢ 1 ▢ 2 ▢ 3 ▢ 4 ▢ 5 ▢ 6 ▢ 7 (Everyday)

---

##### 3. Exhaustion:

- How often does your dog rest (stop, sit, or lie down) during exercise?
  - ▢ Never
  - ▢ Hardly ever
  - ▢ Occasionally
  - ▢ Frequently
  - ▢ Very frequently

---

#### 5. Muscle Condition:

- Forelimb Muscle Condition Score (0 - 3)

  ▢ 0 ▢ 1 ▢ 2 ▢ 3

- Hindlimb Muscle Condition Score (0 - 3)

  ▢ 0 ▢ 1 ▢ 2 ▢ 3

- Epaxial Muscle Condition Score (0 - 3)

  ▢ 0 ▢ 1 ▢ 2 ▢ 3

Sum of MCS:

▢ 0 ▢ 1 ▢ 2 ▢ 3 ▢ 4 ▢ 5 ▢ 6 ▢ 7 ▢ 8 ▢ 9

---

| **Summary of All Frailty Domains**<br>For each domain, if any answer fell within the red zones, that domain receives a score of 1<br>If no answers for a domain fell within the red zones, then that domain receives a score of 0<br>Sum the five domains to determine the **Total Impaired Domains** | | | | |
|---|---|---|---|---|
| NUTRITION | SOCIAL ACTIVITY | EXHAUSTION | MOBILITY | MUSCLE CONDITION |

| **Total Impaired Domains** |
|---|

**Frailty Interpretation**

| Non-Frail | Frail |
|---|---|
| Total Impaired Domains of 0, 1, or 2 | Total Impaired Domains of 3, 4, or 5 |

#### 4. Mobility:

- On average, how frequently does your dog display weakness or lameness in his/her limbs when walking?

  *Weakness/ Lameness is defined as reduced strength in a limb. This can appear as limping (placing less weight on one leg compared to another), increased effort moving the limbs, scuffing of the paws, short strides, or falling.*

  - ▢ Not at all (0 points)
  - ▢ Less than 10% of the time (1 point)
  - ▢ 10 - 50% of the time (2 points)
  - ▢ More than 50% of the time (3 points)

- On average, how frequently does your dog display stiffness in his/her limbs when walking?

  *Stiffness is defined as reduced flexing of the joints. This can present as reduced bending of the limbs, shorter strides, or the dog appearing to be "straight-legged".*

  - ▢ Not at all (0 points)
  - ▢ Less than 10% of the time (1 point)
  - ▢ 10 - 50% of the time (2 points)
  - ▢ More than 50% of the time (3 points)

- Does your dog ever make mistakes while walking?

  *Mistakes may include scuffing of the paws, stumbling, swinging the limbs abnormally outward or inward such that they cross, a lack of coordination between the limbs, and/or skipping steps.*

  ▢ No (0 points) ▢ Yes (1 point)

Sum of Mobility Scores:

▢ 0 ▢ 1 ▢ 2 ▢ 3 ▢ 4 ▢ 5 ▢ 6 ▢ 7

### Supplements and Aging - Changes Since Last Visit

Please complete the survey below. Indicate any changes since your pet's last visit. Thank you!

| | |
|---|---|
| Is this still your current address?<br>[enrollment_arm_1][street_address]<br>[enrollment_arm_1][city_state]<br>[enrollment_arm_1][zipcode] | ◯ Yes ◯ No |
| Please supply your new street address | __________________________ |
| City, State | __________________________ |
| Current zip code | __________________________ |
| Date of address change (approximate) | __________________________ |
| Please describe your home: | ◯ House<br>◯ Duplex<br>◯ Townhouse<br>◯ Apartment<br>◯ Other ______ |
| Which type of area do you and your dog live? | ◯ Rural (arable farming locally)<br>◯ Suburban<br>◯ City/urban |
| Is your dog exposed to pesticides/insecticides/herbicides/ fungicides? (these chemicals are often used in the garden to control insects, plants or fungi) | ◯ Yes ◯ No |
| Please specify type | __________________________ |
| Is this still your preferred phone number?<br>[enrollment_arm_1][phone_number] | ◯ Yes ◯ No |
| Please supply your preferred phone number | __________________________<br>(______) |
| Is this still your preferred email address?<br>[enrollment_arm_1][email1] | ◯ Yes ◯ No |
| Please supply your preferred email address | __________________________ |
| Is [enrollment_arm_1][clinic_name] still [enrollment_arm_1][pet_name]'s primary vet clinic? | ◯ Yes ◯ No |

| | |
|---|---|
| Please provide your current primary vet care provider (clinic name and location). | ________________________________ |
| Are there any changes regarding the other pets living in the household since [enrollment_arm_1][pet_name]'s last recheck on [visit_month_arm_1][visit_date][previous-instance]? Other pets last reported living in the household: [enrollment_arm_1][current_pets_list] | ◯ Yes ◯ No |
| Please specify changes regarding new pets, or pets that are no longer in the household. Please include pet name, and date entered or left the household. For new pets please include species, breed, age, sex, wt. and relationship to [enrollment_arm_1][pet_name] (ex. gets along, dominates, dislikes, ignores, etc.). | ________________________________ |
| Are there any changes to [enrollment_arm_1][pet_name]'s routine medications since [visit_month_arm_1][visit_date][previous-instance]? Current medications listed: [enrollment_arm_1][current_meds_list] | ◯ Yes ◯ No |
| Please indicate medication changes, name, dose, approximate date started or date discontinued. | ________________________________ |
| Have you noticed a change in the usage (dose or frequency) of any medications for pain or anxiety that you give on an as-needed basis since your last visit on [visit_month_arm_1][visit_date][previous-instance]? Current as-needed medications listed: [enrollment_arm_1][prn_meds] | ◯ Yes ◯ No ◯ Not Applicable |
| Has this need increased or decreased? | ◯ Increased ◯ Decreased |
| Please specify medication(s) and how the dose and frequency of any as-needed pain or anxiety drug(s) has changed. | ________________________________ |
| Has [enrollment_arm_1][pet_name] developed any new medical problems since [visit_month_arm_1][visit_date][previous-instance]? Or have any previous medical problems resolved? Ongoing medical problems last indicated: [enrollment_arm_1][current_medical_list] | ◯ Yes ◯ No |
| Please indicate medical problem, date of onset and/or date resolved. | ________________________________ |
| Have you noticed a change in [enrollment_arm_1][pet_name]'s behavior or cognition since your last visit on [visit_month_arm_1][visit_date][previous-instance]? | ◯ Worse ◯ The Same ◯ Improved |

Have you noticed a change in [enrollment_arm_1][pet_name]'s mobility since your last visit on [visit_month_arm_1][visit_date][previous-instance]?

◯ Worse ◯ The Same ◯ Improved

Compared to your last visit on [visit_month_arm_1][visit_date][previous-instance], [enrollment_arm_1][pet_name]'s level of happiness is:

◯ Much worse ◯ Slightly worse ◯ The same ◯ Slightly better ◯ Much better

Have you noticed any of the following signs since [enrollment_arm_1][pet_name]'s last visit on [visit_month_arm_1][visit_date][previous-instance]? (check all that apply)

☐ vomiting
☐ diarrhea
☐ loss of appetite
☐ itching
☐ ataxia/wobbly gait
☐ increased water intake
☐ increased anxiety
☐ changes in behavior, describe ______
☐ other, describe ______
☐ none of the above

#### Adverse Events by System, Severity and Treatment Group

| | | | Placebo | | | Low Dose | | | Full Dose | | |
|---|---|---|---|---|---|---|---|---|---|---|---|
| | | | Month 1 | Month 3 | Month 6 | Month 1 | Month 3 | Month 6 | Month 1 | Month 3 | Month 6 |
| **System Category** | **Adverse Event** | **VCOG Grade** | | | | | | | | | |
| **Allergic/Immunologic Event** | Anaphylaxis | 2 | | | | | | | | | n=1 |
| **Body Cavity** | Ascites | 4 | | | n=1 | | | | | | |
| **Cardiac** | Arrhythmia | 1 | n=1 | | | | | | | | |
| | Hypertension | 2 | | | | | n=1 | | | | |
| | | 1 | | | | n=1 | | | | | |
| **Dermatologic/Skin** | Acute Dermatitis | 2 | | n=1 | | | | | | | |
| | Atopic Dermatitis | 2 | | | | | | | n=1 | n=1 | |
| | Hygroma | 1 | | | | n=1 | | | | | |
| | Preputial infection | 2 | | | | n=1 | | | | | |
| | Pruritus | 1 | n=1 | n=1 | | n=1 | n=4 | n=3 | n=3 | n=2 | n=2 |
| **Dental** | Tooth Decay | 3 | | n=1 | | | | | | | |
| | Tooth Root abscess | 3 | n=1 | | | | | | | | |
| **Ear Disorders** | Ear infection | 2 | | n=1 | | | n=1 | | | | |
| **Gastrointestinal** | Anal Gland Infection | 2 | | | | n=1 | | | | | |
| | Acid Reflux | 1 | | | | | n=1 | | | | |
| | Appetite-Increased | 1 | n=1 | n=2 | n=1 | n=4 | n=2 | n=3 | n=1 | | n=1 |
| | | 2 | n=1 | | n=1 | | n=1 | | | | |
| | Appetite-Decreased | 1 | | n=1 | n=1 | n=2 | n=3 | n=1 | n=3 | n=2 | n=2 |
| | Constipation | 1 | | | | | n=1 | | | | |
| | Diarrhea | 2 | | | | n=1 | n=2 | | | | |
| | | 1 | n=3 | n=1 | | n=3 | n=1 | n=4 | n=3 | n=3 | n=2 |
| | Fecal Incontinence | 5 | | | | | | n=1 | | | |
| | | 2 | | | n=1 | | | | | | |
| | | 1 | | | | | n=1 | | | | |
| | Gastric Dilation-Volvulus | 5 | | | | | | | | n=1 | |
| | Gastroenteritis | 2 | | | | | | | | | n=1 |
| | Irritable Bowel Disease flare-up | 2 | n=1 | n=1 | | | | | | | |
| | Vomiting | 2 | | | n=1 | n=1 | n=1 | | | | |
| | | 1 | n=3 | | n=1 | n=1 | n=1 | n=1 | n=3 | n=1 | n=2 |

Supplementary Figure S7: Adverse Events and Changes in Labwork by System and Group

| | | | | | | | | | | | |
|---|---|---|---|---|---|---|---|---|---|---|---|
| **Hepatobiliary** | Hepatomegaly | 3 | | | n=1 | | | | | | |
| | Non-obstructive Gallbladder Stone | 1 | | | | n=1 | | | | | |
| **Musculoskeletal** | Hindlimb Weakness | 3 | | | | | | | | | n=1 |
| | | 2 | | | | | | | | n=1 | |
| | Lameness | 2 | n=1 | n=1 | | | | | | n=1 | |
| | | 1 | | | | n=2 | | | | | |
| **Neoplasm** | Adrenal Mass | 2 | | | n=1 | | | | | | |
| | Cardiac Mass | 5 | | | | | | n=1 | | | |
| | Lymphoma | 5 | | | | | | n=1 | | | |
| | Oral Mucosal Mass | 2 | | | | n=1 | | | | | |
| | Splenic Mass | 5 | | n=1 | | | | | | | |
| | Soft Tissue Sarcoma | 3 | | | | | n=1 | | | | |
| | Tail Mass | 3 | | | | | | n=1 | | | |
| | Thoracic Mass | 1 | | | n=1 | | | | | | |
| **Neurology** | Acute Neck Pain | 2 | | | | | | n=1 | | | |
| | | 1 | | | | | | | n=1 | | |
| | Anxiety | 1 | n=1 | n=3 | | n=2 | n=1 | n=1 | n=1 | n=2 | n=3 |
| | Ataxia/Wobbly gait | 1 | n=2 | n=3 | n=4 | n=2 | n=2 | n=2 | n=3 | | n=4 |
| | Depression | 1 | | | | n=1 | | | | | |
| | Horner's Syndrome | 1 | | n=1 | | | | | | | |
| | Cauda Equina Syndrome | 3 | | | n=1 | | | | | | |
| | Seizure | 2 | | | | | n=1 | n=1 | | | |
| | | 1 | | | | n=1 | | | | | |
| | Vestibular Episode | 2 | | | | n=1 | | | | n=1 | |
| | | 1 | | | | | | n=1 | | | n=1 |
| **Ocular** | Corneal Ulcer | 2 | n=1 | | | | | | | | n=1 |
| | Refractory Ulcer | 2 | | | n=1 | | | | | | |
| | Ocular Discharge | 1 | | | | n=1 | | | | | n=1 |
| **Renal/Genito** | Bladder Stones | 2 | | | | | | n=1 | | | |
| | Polyuria/ Polydipsia | 2 | | n=2 | n=1 | n=1 | | | | | n=2 |
| | | 1 | n=5 | n=5 | n=5 | n=3 | n=3 | n=3 | | n=1 | |
| | Urinary Incontinence | 5 | | | | | | n=1 | | | |
| | | 1 | | | | n=1 | n=1 | | | | |
| | Inappropriate Urination | 1 | | n=2 | n=3 | n=2 | n=2 | n=1 | n=2 | n=1 | n=3 |
| **Resp/Pulm** | Aspiration Pneumonia | 2 | | | | | | n=1 | | | |
| | Coughing | 1 | | | | n=1 | n=1 | | | n=1 | |

Supplementary Figure S7: Adverse Events and Changes in Labwork by System and Group

| | | | | | | | | | | | |
|---|---|---|---|---|---|---|---|---|---|---|---|
| | Collapsed Lung/ Pneumonia | 5 | | | n=1 | | | | | | |
| | Increased Respiratory Effort | 1 | n=1 | | | | | | | | |
| | Panting | 1 | | | | n=2 | | n=1 | | | n=1 |
| | Tracheal Collapse | 2 | | | | | | n=1 | | | |

#### Lab Work Changes by System, Severity and Treatment Group

| | | | **Placebo** | | | **Half Dose** | | | **Full Dose** | | |
|---|---|---|---|---|---|---|---|---|---|---|---|
| | | | **Month 1** | **Month 3** | **Month 6** | **Month 1** | **Month 3** | **Month 6** | **Month 1** | **Month 3** | **Month 6** |
| **System Category** | **Adverse Event** | **VCOG Grade** | | | | | | | | | |
| **Blood/Bone Marrow** | Anemia | 1 | n=2 | | n=5 | n=6 | | n=4 | n=1 | | n=3 |
| | Lymphopenia | 1 | n=4 | | n=1 | n=5 | | n=4 | n=2 | | n=5 |
| | Monocytosis | 1 | n=2 | | | n=1 | | | n=1 | | |
| | Monocytopenia | ` | | | | | | | n=1 | | |
| | Plasma Protein-Decreased | 1 | n=1 | | n=2 | | | | n=1 | | n=1 |
| | Plasma Protein-Increased | 1 | n=4 | | n=1 | n=3 | | n=1 | n=1 | | |
| | Stress Leukogram | 1 | | | n=1 | n=1 | | n=1 | | | |
| | Thrombocytosis | 1 | | | n=2 | | | | | | |
| | Thrombocytopenia | 1 | | | | | | | n=1 | | n=1 |
| **Metabolic/ Laboratory** | Albumin-Increased | 1 | | | | | | n=1 | | | |
| | Amylase-Increased | 1 | | | n=2 | | | n=1 | | | n=1 |
| | Anion Gap-Decreased | 1 | | | | | | | | | n=1 |
| | ALP-Increased | 1 | n=1 | | n=2 | n=3 | | n=3 | | | n=1 |
| | ALT-Increased | 3 | | | | | | | | | n=1 |
| | | 1 | | | n=1 | | | n=3 | | | n=2 |
| | AST-Increased | 1 | | | n=1 | n=2 | | n=1 | n=2 | | n=1 |
| | Bicarbonate-Decreased | 1 | n=1 | | n=1 | | | | | | |
| | Bicarbonate-Increased | 1 | n=1 | | | | | | n=1 | | |
| | BUN-Decreased | 1 | n=1 | | n=1 | | | | | | |
| | BUN-Increased | 2 | | | | | | n=1 | | | n=1 |
| | | 1 | n=1 | | | n=1 | | n=1 | n=3 | | n=1 |
| | Creatinine-Decreased | 1 | | | n=1 | n=1 | | | | | |
| | Creatinine-Increased | 1 | | | | n=2 | | n=1 | n=1 | | n=2 |

Supplementary Figure S7: Adverse Events and Changes in Labwork by System and Group

| | | | | | | | | | | | |
|---|---|---|---|---|---|---|---|---|---|---|---|
| | Creatine Kinase-Increased | 1 | n=2 | | | n=4 | | n=1 | | | |
| | GGT-Increased | 1 | | | n=1 | n=1 | | n=1 | | | n=3 |
| | Globulins-Decreased | 1 | n=1 | | n=2 | | | | | | n=1 |
| | Hyperkalemia | 1 | n=3 | | n=1 | n=1 | | | | | n=1 |
| | Hypercalcemia | 1 | | | | | | n=2 | n=1 | | n=4 |
| | Hyperchloremia | 1 | | | n=1 | | | | | | |
| | Hypermagnesemia | 1 | | | | | | | n=1 | | n=2 |
| | Hyperphosphatemia | 1 | | | n=1 | | | | n=1 | | n=1 |
| | Hypocalcemia | 1 | | | n=1 | n=1 | | | | | |
| | Hypochloremia | 1 | | | n=1 | n=1 | | | n=1 | | |
| | Hypoglycemia | 1 | | | | | | | | | n=1 |
| | Hypomagnesemia | 1 | n=1 | | n=1 | n=1 | | n=1 | n=1 | | |
| | Hyponatremia | 1 | n=2 | | n=5 | n=1 | | n=1 | n=3 | | n=3 |
| | Hypophosphatemia | 1 | | | | n=2 | | n=2 | n=1 | | n=1 |
| | Lipase-Increased | 1 | n=1 | | n=3 | n=1 | | | | | n=2 |
| | Total Protein-Decreased | 1 | n=1 | | n=1 | n=1 | | | | | |
| | Bacteriuria | 1 | n=1 | | n=3 | n=1 | n=1 | n=1 | n=4 | n=1 | n=2 |
| | Bilirubinuria | 1 | | | n=3 | n=1 | n=1 | | n=1 | n=1 | n=1 |
| **Renal/ Genitourinary** | Hematuria | 1 | n=3 | n=1 | n=5 | n=5 | n=2 | n=4 | n=4 | | n=2 |
| | Isosthenuria | 1 | | | n=1 | n=1 | | | | | n=1 |
| | Ketonuria | 1 | n=3 | | n=6 | n=3 | n=1 | n=3 | n=3 | | n=1 |
| | Proteinuria | 1 | n=1 | | | n=5 | n=1 | n=3 | n=1 | | n=1 |

Supplementary Table S7: Adverse Events and Labwork

Supplementary Figure S7: Adverse Events and Changes in Labwork by System and Group

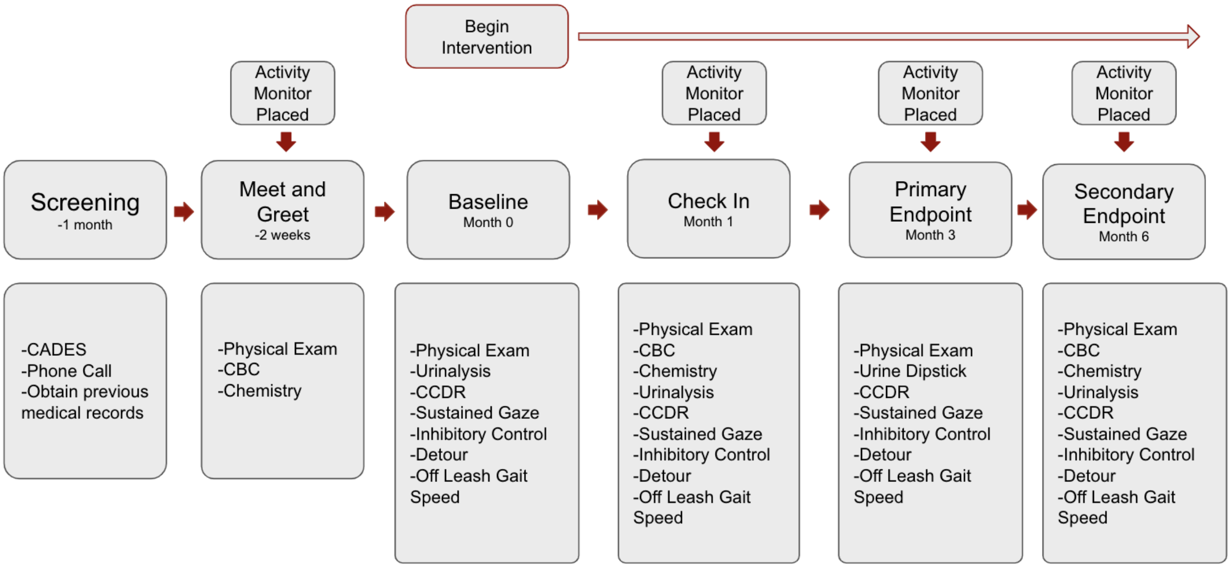

Supplementary Figure S8: Study Timeline. The timeline of all visits in the study are provided along with all assessments performed at each visit. Activity monitors were placed at specific visits in order to obtain two week increments of activity data for each timepoint. Intervention was started the day following an individual's baseline (month zero) visit.
